# PolyA tail segmentation improves the stability of the template DNA and increases the translatability of *in vitro* transcribed mRNA

**DOI:** 10.1101/2025.10.21.683660

**Authors:** Tomasz Spiewla, Karol Czubak, Zofia Pilch, Marek R. Baranowski, Pawel S. Krawczyk, Kamila Affek, Wiktor Antczak, Marta Szulc-Gasiorowska, Sebastian Chmielinski, Seweryn Mroczek, Michal Brouze, Dominika Nowis, Jakub Golab, Andrzej Dziembowski, Jacek Jemielity, Joanna Kowalska

## Abstract

PolyA tail regulates mRNA localization, stability, and translation. PolyA length affects the durability and translational activity of both endogenous and exogenously delivered mRNAs. However, long polyA stretches can undergo recombination during amplification in bacterial plasmids, impairing the production of *in vitro* transcribed (IVT) mRNA with long polyAs. PolyA tail segmentation with heteronucleotide spacers has recently emerged as a solution. Here, we developed segmented polyA patterns that stabilize the sequence during DNA amplification and enhance mRNA translation. We designed 15 novel genetically modified polyA variants, differing in the length, placement, and frequency of spacers, and the overall length (from ∼120 to ∼200 nucleotides). We evaluated their stability in DNA plasmids and homogeneity, translational activity, and durability in cell culture of the resulting mRNAs, comparing them to A_90_ tail and other known solutions, including those from existing mRNA vaccines. Selected sequences were validated *in vivo*. Surprisingly, we found that even frequent heteronucleotide insertions produce functional polyA tails. The most notable enhancements in protein production were observed for a segmented tail exceeding 200 nt in length [A_30_(CA_15_)_11_; up to 6-fold compared to mRNA with A_90_ tails]. Our findings extend the scope of possible polyA modification strategies, offering new possibilities for advancing mRNA therapeutics.

## INTRODUCTION

The efficacy of mRNA-based therapeutics relies heavily on the structural properties of the mRNA molecule, which govern both its stability and translation efficiency. A typical mRNA consists of a coding sequence flanked by untranslated regions (UTRs) that regulate its expression, along with structural elements at both termini that enhance its translation efficiency and molecular stability. The 5’-end is capped with 7-methylguanosine (m^7^G), while the 3’-end terminates in a polyA tail, a homopolymer sequence of adenine nucleotides of varying length.(1) The polyA tail of eukaryotic messenger RNAs (mRNAs) is a highly dynamic element that undergoes significant changes throughout the mRNA lifecycle. Once transcription has ended and primary polyadenylation has occurred, the polyA tail is usually between 150 and 250 adenines long. This initial polyadenylation is carried out by the canonical polyA polymerase (PAP), whose activity is stimulated and regulated by nuclear polyA-binding proteins known as PABPNs.(2–4) After export of mRNA to the cytoplasm, polyA tails are recognized by cytoplasmic polyA-binding protein (PABPC), which facilitates the formation of the translation initiation complex (5) and the recruitment of ribosomes, as well as prevents mRNA backtracking to the nucleus.(6) The polyA tails undergo dynamic shortening, mainly by the CCR4-NOT complex, and some secondary elongation by non-canonical polyA/U polymerases, leading to a steady-state length of approximately 80–100 nt in mammalian cells.(7) This variability plays a crucial role in regulating mRNA stability and translational efficiency. Bulk mRNA degradation pathways begin with the mRNA deadenylation, i.e. enzymatically catalysed shortening of the polyA.(8) A minimum polyA tail length of around 30 nucleotides (nt) is usually required to guarantee the stability of mRNA and its efficient translation.(9) This is because it corresponds to the binding footprint of polyA-binding proteins (PABPs), which protect mRNA from degradation.(9) Polyadenylation dynamics influences mRNA half-life, directly correlating polyA tail length with transcript stability and protein production.(8,10) Long polyA tails are typically associated with enhanced translation and prolonged mRNA lifespan, whereas shortened polyA tails signal mRNA degradation, reducing its half-life and protein output.(10,11) PolyA tail length also significantly affects the protein output from exogenously delivered mRNAs.(2,12–14) Recently, it has been demonstrated that antigen-encoding mRNAs can undergo TENT5A-mediated polyadenylation in the cytoplasm, which enhances their therapeutic efficacy.(15) In view of this critical regulatory role, strategies to stabilize and optimize polyA tails in mRNA have become an increasingly appealing approach to designing mRNA-based therapeutics.

The polyA tail has become an increasingly frequent target for chemical and chemoenzymatic modifications, advancing mRNA-based technologies.(12,16–21). These include branched and topologically modified, so-called multimeric, tails or introducing chemical groups such as 2′-O-methyl (2MOE) or terminal dideoxyctidines ddC to prevent degradation. (22) RNA circularization has also been explored as an alternative to polyA modification.(23–25) Overall, unmodified polyA tails can be incorporated into IVT mRNA using two methods. The first approach, commonly used in the manufacturing of mRNA-based therapeutics, relies on encoding the polyA tail in the DNA template sequence, allowing for the straightforward generation of polyadenylated mRNA during *in vitro* transcription. This approach, in principle, enables the most precise control of polyA tail length and homogeneity. However, long polyA stretches undergo recombination in *E. coli* strains commonly used for DNA plasmid amplification, resulting in the shortening of polyA sequences and an increase in heterogeneity. The alternative method relies on the generation of polyA tails by template-independent polymerase, such as bacterial polyA polymerase.(26,27) This approach enables the generation of longer tails and is more compatible with certain chemical modifications, but the length and homogeneity of the polyA are challenging to control. Collectively, these strategies provide a modular and adaptable toolbox for optimizing synthetic mRNA molecules; however, only the template-encoded approach is fully compatible with existing mRNA manufacturing pipelines.

Using the template-encoded approach, mRNAs with polyA tails no longer than 90–120 nucleotides can be reliably generated. To address length-related limitations, researchers have developed a strategy to stabilize polyA tract during plasmid amplification by dividing it into two or more polyA segments separated by a heteronucleotide spacer. This approach has been employed in the design of the BNT162b2 COVID-19 mRNA vaccine developed by Pfizer/BioNTech, which contains 30-nt polyA tract followed by a 10-nt spacer and a 70-nt poly(A) tract.(28) Conversely, the terminal pentamer sequence (mΨCmΨAG) at the 3′ end of Moderna’s mRNA-1273 vaccine, which remains after plasmid linearisation before *in vitro* transcription (IVT), serves as a specific sequence for easily discriminating intact vaccine mRNA molecules. Its physiological function may delay the deadenylation process, thereby enhancing stability.(15) To our knowledge, the performance of these two polyA sequences has never been directly compared. Trepotec et al. found that dividing a 120-adenine tract into two or three equal parts by mono-or hexanucleotide spacers reduces recombination events in plasmid DNA with no adverse effects on mRNA stability or translation.(29) Interestingly, Li et al. (2022) modified the 40-nt polyA tail by incorporating short heteronucleotide sequences. They revealed that cytidine (C) residues near the polyA tail terminus enhance protein production in multiple cell lines (30).

Intrigued by these previous findings, we decided to further explore the potential of polyA tail segmentation to achieve two independent goals: (i) mitigate the instability of long homopolymeric sequences in DNA templates encoding mRNA and (ii) enhance the translational efficiency of mRNA. We investigated mRNAs with polyA homosequence replaced by 15 various novel heterosequences, consisting of adenine stretches of different lengths interspersed with mono-or oligonucleotide spacers. Importantly, we have compared these modified tails to the previously developed segmentation patterns, including the ones present in approved mRNA vaccines. We identified sequence motifs that increase protein output in multiple tested cell lines and *in vivo* (mouse models) up to 10-fold compared to mRNA with unmodified A_90_ polyA. We determined how these modifications affect the durability and translational activity of mRNA, shedding new light on mRNA structure-activity relationship. Hence, our study presents a robust framework for optimizing polyA tail modifications, with direct implications for mRNA-based therapeutics and gene expression technologies.

## MATERIALS AND METHODS

### Preparation of DNA inserts with designed 3’-tail modifications

Plasmid DNA vectors with modified 3’-tail sequences were constructed using double-stranded DNA oligonucleotide inserts (Table S1). Each insert was designed and ligated into the pJet1.2 or pUC57 vector encoding *Firefly* luciferase, mKate2-PEST, human erythropoietin (hEPO), and human alpha-1-antitripsin (hA1AT) genes using a blunt-end cloning approach. For insert preparation, complementary DNA oligonucleotides (Metabion) were mixed in equimolar ratios (final 60 µM per strand) and phosphorylated using T4 Polynucleotide Kinase (NEB) in T4 PNK buffer (NEB) at 37°C for 30 min. Hybridization was performed by heating to 95°C, followed by gradual cooling to 25°C over 2 h (gradient ∼2/3 °C/min). Inserts were purified using a commercial DNA purification kit (Macherey-Nagel) according to the manufacturer’s protocol. Each experimental variant (R^1^–R^6^, S^1^–S^15^) was prepared following this protocol, with specific sequences listed in Table S2.

### Preparation of plasmid vectors

The circular plasmid encoding *Firefly* luciferase, mKate2-PEST, hEPO, or hA1AT (5 µg) was linearized with AarI restriction enzyme (Thermo) in 10× AarI buffer (Thermo) and 50× Oligo (Thermo) at 37°C for 16 h. The linearized plasmid was then purified using a commercial DNA purification kit (Macherey-Nagel). To generate blunt ends, the enzymatic reaction was performed using DNA Polymerase I, Large Fragment (Klenow) in 10× Buffer 3 (NEB) with 10 mM dNTP (Thermo) at 25°C for 15 min, followed by another purification step. The plasmid was then dephosphorylated using FastAP Thermosensitive Alkaline Phosphatase (Thermo) in 10× FastAP buffer (Thermo) at 37°C for 10 min, and the final purified vector (V1) was obtained.

### Ligation and bacterial transformation

A total of 100 ng of the linearized vector (V1) was mixed with DNA inserts (R^1^–R^6^, S^1^–S^15^) at a molar ratio of 1:3 (vector:insert). The ligation reaction was carried out by adding 10× Ligase buffer (Thermo), PEG 2000 (Thermo), ATP (Thermo), and T4 DNA Ligase (Thermo) and incubating at 25°C for 1 h. The reaction mix (20 µL) was then cooled to 4°C and incubated for 1 h. For bacterial transformation, 10 µL of the ligation mixture was added to 50 µL of pre-chilled chemically competent bacteria (NEB Stable Competent *E. coli*) and incubated on ice for 30 min. Heat shock transformation was performed at 42°C for 30 s, followed by cooling on ice for 2 min. 500 µL of NEB Stable outgrowth medium (NEB) was then added, and the mixture was incubated at 30°C for 1 h with shaking (300 RPM). Transformed cells (200 µL) were plated on LB-agar (Roth) supplemented with 100 µg/mL Ampicillin (Roth) and incubated at 30°C for 16 h. Single colonies corresponding to each variant (R^1^–R^6^, S^1^–S^15^) were selected and inoculated in 5 mL of LB medium (Roth) with 100 µg/mL Ampicillin, followed by incubation at 30°C for 16 h.

### Plasmid purification and validation

Bacterial cultures were centrifuged (4000×g, 10 min), and plasmid DNA was purified using the GeneJET Plasmid Miniprep Kit (Thermo). Plasmid concentration was measured using a NanoDrop 2000c spectrophotometer (Thermo), and sequencing was performed using the Sanger method (Genomed, Poland) to confirm the incorporation of modified 3’-tail sequences. Once confirmed, fresh transformations were performed following the previous protocol. A single colony was selected, inoculated in 200 mL of LB medium (Roth) with 100 µg/mL Ampicillin, and incubated at 30°C for 16 h. Large-scale plasmid purification for all variants (R^1^– R^6^, S^1^–S^15^) was performed using the E.Z.N.A.^®^ FastFilter Plasmid DNA Maxi Kit (Omega).

### Plasmid linearization and template preparation for *in vitro* transcription

Plasmid vectors encoding *Firefly* luciferase, mKate2-PEST, hEPO, or hA1AT with modified 3’-tail sequences (R^1^–R^6^, S^1^–S^15^) were linearized by incubating 30 µg of plasmid DNA with AarI enzyme (Thermo), 10× AarI buffer (Thermo), and 50× Oligo (Thermo) at 37°C for 16 h. The linearized plasmid templates were purified using a commercial DNA purification kit (Macherey-Nagel). To verify template integrity, approximately 100 ng of DNA from each variant (R^1^–R^6^, S^1^–S^15^) was loaded onto a 1% agarose gel (VWR) with 6× loading dye (Thermo). Electrophoresis was performed in 1× TAE buffer at 140 V for 25 min.

### Instability analysis of polyA tails in plasmid DNA vectors

Chemocompetent *E. coli* Top10 and NEB^®^ Stable Competent *E. coli* (High Efficiency) were transformed with plasmids carrying modified polyA (S^1^–S^15^) and reference (R^1^–R^6^) sequences, according to the procedure described above. For each variant, 30 individual colonies were selected from bacterial plates, cultured in liquid medium, and subjected to plasmid purification. The purified plasmids were then sequenced by the Sanger method to assess polyA tail length and sequence integrity after amplification. The number of clones displaying deviations in polyA tail length and uniformity was recorded, and the proportion of bacterial colonies exhibiting shortened polyA sequences after plasmid replication was calculated and reported.

### *In vitro* transcription and purification of mRNA

*In vitro* transcription (IVT) of FLuc, mKate2_PEST, hEPO, and hA1AT mRNA with modified 3’-tail sequences (R^1^–R^6^, S^1^–S^15^) was performed using linearized DNA template containing the T7 promoter Φ6.5 (TAATACGACTCACTATAGGG), transcription buffer (10×, Thermo), GTP (4.0 mM, Thermo), CTP (5.0 mM, Thermo), ATP (5.0 mM, Thermo), m^1^Ψ-UTP (5.0 mM, Jena Bioscience), m^7^GpppA_m_pG-cap1 (10 mM) (31), MgCl_2_ (15 mM), Ribolock (1U/µL, Thermo), Inorganic Pyrophosphatase (0.002U/µL, Thermo), T7 RNA Polymerase (0.125 mg/mL). After incubation for 1 h at 37°C, DNase I (6 µL, 30 min, Thermo) was added, and incubation continued for another 30 min. The mixture was then diluted twice with water, and an EDTA solution (9 µL, 500 mM, VWR) was added to inhibit the reaction. The crude RNA for all variants (R^1^–R^6^, S^1^–S^15^) was purified after IVT using POROS™ Oligo (dT)_25_ Affinity Resin (Thermo) by applying the entire reaction volume to a previously conditioned resin (2 mL) of Oligo (dT)_25_ (20mM Tris pH 7.4, 800 mM NaCl) and incubated (10 min, 25°C). The resin was then washed with equilibration buffer (5 volumes, 20 mM Tris pH 7.4, 800 mM NaCl) and wash buffer (5 volumes, 20 mM Tris pH 7.4, 300 mM NaCl). The elution was performed by incubating the resin with water (2 times 3 volumes each, 5 min, 65°C). RNA was filtered (0.22 µM), concentrated (Amicon 100 kDa, 5000×g, 4°C, 15 min), and further purified via HPLC (RNASep™ Prep, ADC Biotec) using a TEAA-MeCN gradient Buffers A: 100 mM TEAA, B: 100 mM TEAA, 100 mM MeCN. Program: 20% B for 5 min, 20-29% B in 20 min, 29-100% B in 1 min, 100% B for 4 min, flow 5.0 mL/min, 55°C (RT ∼ 35 min). RNA integrity was verified by agarose gel electrophoresis (1%, 1× TBE, 140 V, 20 min), with ∼100 ng of each RNA variant (R^1^–R^6^, S^1^–S^15^) applied. Dot-blot analysis was performed to detect double-stranded RNA (dsRNA), using 25 and 250 ng of each variant applied to a nylon membrane (Sigma), followed by J2 antibody (SCICONS) and HRP-conjugated secondary antibody (Thermo) detection. A dsRNA ladder (NEB, 25 ng) served as a positive control. Signals were visualized using chemiluminescence (Immobilon Western, Amersham Imager 600, Cytiva).

### Nanopore sequencing

#### DNA

Plasmids encoding mRNAs with polyA tails variants R^1^, S^14^ and S^15^ were subjected to the nanopore sequencing using MinION device (ONT), flowcell FLO-MIN1114 and sequencing kit SQK-RBK114-24. Following sequencing, reads were basecalled using dorado (v 1.1.1, ONT), with polyA length determination turned on (--estimate-poly-a) and sequence-specific settings for polyA localization in raw nanopore signal. After demultiplexing and mapping to the reference sequences, histograms of polyA lengths were generated for reads covering the whole reference sequence, and with mapping quality>0.

#### RNA

Direct RNA sequencing (DRS) was performed as described by Bilska.(32) The 50-100 ng of IVT mRNA was used for library preparation with a Direct RNA Sequencing Kit (catalog no. SQK-RNA002, Oxford Nanopore Technologies) according to the manufacturer’s instructions. Sequencing was performed using R9.4 flow cells on a MinION device (ONT). Raw data were base called using Guppy(ONT). Raw sequencing data (fast5 files) were deposited at the European Nucleotide Archive (ENA, accession numbers to be provided).

### Cell culture and transfection

A549 (human lung carcinoma, ATCC CCL-185), HEK293T (human kidney epithelium, ATCC CRL-3216), and HepG2 (human hepatocellular carcinoma, ATCC HB-8065) cells were cultured in DMEM (Gibco) supplemented with 10% FBS (Sigma), GlutaMAX (Gibco), and 1% penicillin/streptomycin (Gibco) at 37°C, 5% CO₂. JAWS II (mouse dendritic cells, ATCC CRL-11904) were maintained in RPMI 1640 (Gibco) with 10% FBS, sodium pyruvate (Gibco), 1% penicillin/streptomycin, and 5 ng/mL GM-CSF (PeproTech) under the same conditions. One day before the transfection, cells were seeded in 96-well plates (1×10⁴ cells/well). The next day, RNA constructs with different polyA tail variants were transfected in triplicate per cell line using Lipofectamine MessengerMAX (Invitrogen) according to the manufacturer’s protocol. Each reaction was prepared by pre-incubating 0.15 μL Lipofectamine in 5 μL Opti-MEM (Gibco) for 10 min at room temperature (Mix 1). In parallel, 50 ng RNA was diluted in 5 μL Opti-MEM (Mix 2). Mix 1 and Mix 2 were combined, incubated for 5 min, and added to the cells.

### *In vitro* FLuc expression analysis

Following transfection, cells were incubated at 37°C, 5% CO₂. Luciferase activity was measured at 4 h, 16 h, 24 h, 48 h, and 72 h post-transfection using the Bright-Glo Luciferase Assay System (Promega) following the manufacturer’s instructions. Luminescence was recorded in white, opaque 96-well plates using an EnVision plate reader (Perkin Elmer, Waltham, MA, USA).

### *In vitro* mKate2_PEST expression analysis

HEK293T and A549 cells used in transfection were cultured in DMEM medium (Thermo Fisher Scientific) supplemented with 10% FBS (Gibco) at 37°C in 5% CO_2_. Reverse transfection was performed in a 384-well plate (Perkin Elmer) using cells 9 days after thawing (1.4×10^3^cells/well for HEK293T and 1.6×10^3^cells/well for A549). Transfection complexes were pre-deposited onto the wells prior to cell seeding. Transfection of each tested mRNA was prepared in three technical replicates. The mRNA used for transfection was diluted in Opti-MEM (Gibco) in a 96-well plate (Greiner). The mixture was prepared to obtain 40ng of mRNA in 10μL of Opti-MEM and dispensed onto a 384-well plate. A dilution of MessengerMAX (Invitrogen) in Opti-MEM equivalent to 0.1μL of reagent in 10μL of Opti-MEM per well of the 384-well plate was prepared in a 5mL tube. As with mRNA, 10μL of MessengerMAX dilution was dispensed onto a 384-well plate. During a 40 minutes incubation, cells were detached and counted in a Countess 3L cell counter (Invitrogen) using Countess cell counting chamber slides (Invitrogen). Two suspensions of each cell line in a volume of 100μL were seeded on the 384-well plate using Multipette E3x (Eppendorf). Non-confocal imaging was started 5 hours after transfection in Opera Phenix PerkinElmer using a water objective 20x in three channels - brightfield, DPC and mCherry. During 72-hours imaging, cells were maintained under standard conditions of 37 °C and 5% CO₂. Pictures were captured every two hours. Primary image analysis including cell segmentation was conducted in Harmony 4.9 software (PerkinElmer). Cells were segmented based on Digital Phase Contrast. For each segmented cell mean intensity values from technical triplicates were calculated. Exported values were processed in R (version 4.2.2).

### Statistical analysis

Data points represent biological replicates. Bars represent means ± standard deviation (SD) or medians (where stated). If data passed normality test using Shapiro-Wilk method, one way ANOVA with Tukey’s multiple comparisons test was used. If data did not pass normality test using Shapiro-Wilk method, Kruskal-Wallis test with Dunn’s multiple comparisons test was used. For all analyses, differences between the groups were considered statistically significant at P<0.05. P values and statistical test used are provided in the figure captions. Statistical analysis was performed in Prism version 9.5.1 (GraphPad Software Inc., San Diego, CA).

### Model fitting of mRNA and protein dynamics

The fluorescence data were analysed using a system of differential equations to model the dynamics of mRNA and protein levels. The model included parameters for mRNA decay, translation rate, protein maturation, and protein decay, as described in Mauger *et al*. (33): d[mRNA] = -[mRNA] λ RNA, d[inactive protein] = [mRNA] k trans - [inactive protein] k mat, d[fluor] = [inactive protein] k mat – [fluor] λ fluor, where λ RNA is the decay rate of a given mRNA, k trans is the speed of inactive fluorescent peptide production, k mat is the speed of peptide maturation into fluorescently active protein, and λ fluor is the decay rate of the fluorescent protein. The curve_fit function from the scipy.optimize module was used to fit the model to the experimental fluorescent data. Multiple initial guesses were tested to avoid local minima. For each experimental group, the globally optimal parameters for protein maturation and decay were determined using minimize function from the scipy.optimize module, using MAE (Mean Absolute Error) to measure the quality of each fit. This optimization was performed across all subsets of the data to ensure consistent parameter values for protein maturation and decay across different experimental conditions (i.e. cell type). Within each experimental group, the locally optimal parameters for mRNA half-life (t_0.5_ RNA = ln(2) / λ RNA) and protein production rate were calculated for each added reporter mRNA, using the globally optimal parameters for protein maturation and decay. The obtained parameters were averaged across technical replicates and standard deviation values for protein production and half-life were calculated.

### BMDM culture and transfection

Bone marrow-derived macrophages (BMDMs) were obtained from wild-type C57BL/6 adult mice (12–25 weeks old). Mice were sacrificed by cervical dislocation, and femurs and tibias were isolated. Bone marrow was harvested using a centrifugation-based protocol and plated in IMDM medium (Thermo Fisher) supplemented with 10% FBS (Gibco), 100 U/mL penicillin/0.1 mg/mL streptomycin (Sigma-Aldrich), and 10 ng/mL M-CSF (Preprotech). Cells were cultured at 37°C, 5% CO₂ and used for experiments after 14 days of differentiation. Differentiated BMDMs (0.5×10⁶ cells/well) were seeded in 6-well plates one day before transfection. Cells were transfected with 1 µg of IVT mRNA encoding Firefly luciferase (FLuc) using Lipofectamine™ MessengerMAX™ (Invitrogen) according to the manufacturer’s protocol. Cells were harvested 24 h post-transfection for Western blot analyses.

### Western blot analysis of protein expression from mRNA

Cells were lysed in PBS containing 0.1% NP-40, protease inhibitors, and 0.1 U/mL Viscolase (A&A Biotechnology) for 30 min at 37°C with 1200 rpm shaking. Then, 3× SDS sample buffer (187.5 mM Tris-HCl, pH 6.8, 6% SDS, 150 mM DTT, 0.02% bromophenol blue, 30% glycerol, 3% 2-mercaptoethanol) was added, and samples were boiled for 10 min. Proteins were separated on 12–15% SDS-PAGE gels and transferred onto Protran nitrocellulose membranes (GE Healthcare) using wet transfer (400 mA, 4°C, 1.5 h) in 1× transfer buffer (25 mM Tris, 192 mM glycine, 20% methanol). Membranes were stained with 0.3% Ponceau S in 3% acetic acid, digitized, and blocked in 5% milk in TBST for 1 h. Primary antibodies were incubated overnight at 4°C at the following dilutions: 1:3000 (mKate2, Firefly luciferase), 1:5000 (actin, tubulin) in 5% milk in TBST. Membranes were washed 3× in TBST (10 min each) and incubated with HRP-conjugated secondary antibodies: anti-mouse (Millipore, 401215) and anti-rabbit (Millipore, 401393), both at 1:5000, for 2 h at RT. After 3× washes in TBST, protein detection was performed using the ChemiDoc System.

### mRNA formulation into LNPs

The SM-102 lipid was purchased from BroadPharm (cat. number BP-25499, USA). The 1,2-dimyristoyl-rac-glycero-3-methoxypolyethylene glycol-2000 (DMG-PEG2k), cholesterol (ovine wool) and 1,2-dioctadecanoyl-sn-glycero-3-phosphocholine (DSPC) were purchased from Avanti Polar Lipids (USA). The stock solutions of lipids were prepared in absolute ethanol (Thermo Fisher Scientific) at a concentration of 100 mg/mL for SM-102 and 10 mg/mL for the rest of the lipids. The lipid mix was prepared by combining SM-102, DSPC, cholesterol and DMG-PEG2k at a molar ratio of 50:10:38.5:1.5 in absolute ethanol at a total concentration of 12.5 mM. Stock solution of mRNA was diluted in 100 mM sodium citrate buffer pH 4.0 at the mRNA final concentration of 114 ng/µL. The lipid mix and mRNA solution were combined in a microfluidic device (NanoAssemblr^®^ IgniteTM, Precision NanoSystems) equipped with NxGen Cartridge at a flow ratio of 3:1 (aqueous phase:ethanol) with a total flow rate of 12 mL/min. The final weight ratio of SM 102:mRNA was 13:1 and N/P ratio (cationic nitrogen groups from the ionizable lipid over anionic phosphate groups from the mRNA) was ∼6. All resulting LNP nanoparticles (LNPs) were diluted 20-40 times in 1×PBS sterile buffer (w/o calcium and magnesium ions) and concentrated by ultrafiltration using Amicon ultracentrifugal tubes (Merck Millipore) with 100 kDa MWCO (2500xg, 20°C). LNPs were stored at 4°C for no longer than 2 weeks before use. The mRNA concentration in the formulated LNPs was measured using the RiboGreen assay (Thermo), followed by dynamic light scattering (DLS) analysis to determine nanoparticle size (nm) and homogeneity, assessed through the polydispersity index (PDI).

### *In vivo* FLuc evaluation

Experiments were conducted in 10–12-week-old (∼25 g) female BALB/c mice under a protocol approved by the II Local Ethical Committee for Experiments on Animals in Warsaw, Poland (WAW2/126/2021), following Directive 2010/63/EU on animal protection. Mice were maintained under specific pathogen-free (SPF) conditions in individually ventilated cages (IVC) with a 12-hour light/dark cycle and unrestricted access to food and water. Mice (n = 5 per group) received 100 µL of SM-102-encapsulated FLuc mRNA (10 µg per mouse) intravenously. Bioluminescence imaging was performed at 4, 8, 12, and 24 h post-injection using the *In Vivo* Imaging System (IVIS, PerkinElmer). Prior to imaging, mice were anesthetized with isoflurane, and D-luciferin (150 mg/kg, Syd Labs) was administered intraperitoneally 5 minutes before measurement. Bioluminescence was quantified as total photon flux (photons/second) using Living Image software (PerkinElmer).

### *In vivo* hEPO evaluation

Experiments were conducted in 10–12-week-old (∼25 g) female C57BL/6 mice under a protocol approved by the II Local Ethical Committee for Experiments on Animals in Warsaw, Poland (WAW2/085/2023), in accordance with Directive 2010/63/EU. Mice (n = 5 per group) received 100 µL of SM-102-encapsulated hEPO mRNA (1 µg per mouse) intravenously. Blood samples were collected from the submandibular vein at 4 and 24 hours post-injection. Serum was separated by centrifugation (2000×g, 15 min, RT) after clotting at room temperature. Human EPO levels were quantified using an ELISA kit (Invitrogen, #BMS2035-2) following the manufacturer’s instructions.

### *In vivo* hA1AT evaluation

The experiment was conducted on 10–12-week-old (∼25 g) female C57BL/6 mice under a protocol approved by the II Local Ethical Committee for Animal Experiments in Warsaw, Poland. Mice (n = 5 per group) received 100 µL of SM-102-encapsulated hA1AT mRNA (1 µg per mouse) via intravenous injection. Blood samples were collected from the submandibular vein at 4, 24, and 48 hours post-injection. After clotting at room temperature, serum was separated by centrifugation (2000×g, 15 min, RT). Human A1AT levels were quantified using an ELISA kit (Innovative Research, #IHUAATKT) according to the manufacturer’s instructions.

## RESULTS AND DISCUSSION

### Structural polyA design for enhanced DNA template stability and mRNA translation

The therapeutic efficacy of messenger RNA (mRNA) molecules depends on structural features that determine their stability, translational activity, and regulate cellular metabolic processing. In this context, one of the most critical structural elements of mRNA is the polyA tail. The traditional approach, based on long template-encoded homopolymeric adenine sequences at the 3’ end of mRNA, has limitations, including the instability of corresponding plasmids and the consequent maximal polyA tail length.(34) Developing structural mRNA modifications that enhance DNA template stability and improve protein output in living organisms is thus highly desirable. To address this, we first designed eight distinct 3’-tail sequences (S^1^–S^8^) that could potentially replace unmodified polyA tails. In these constructs, we utilized 3’-tail modifications incorporating various heteronucleotide spacers, combined with a fixed segment of 30 consecutive adenines at the 5’ end of the polyA region (5’ A_30_ motif). We intentionally left the 5’-terminal segment unmodified, assuming that it can be crucial for effective binding of the first PABP molecule, which recognizes approximately 27 nucleotides of the polyadenylated chain.(35–38) We compared these with reference constructs representing the current state-of-the-art, which included unmodified polyA, lead structures from previous literature reports, and polyA variants used in approved mRNA-based vaccines. Based on the initial results, which indicated that even densely segmented polyA tails remain translationally active, we later designed seven additional sequences (S^9^–S^15^) to further optimize and better understand the structure-activity relationship. We began by constructing DNA oligonucleotide sequences (Table S1), which were subsequently incorporated into plasmid DNA vectors pJet and pUC57 encoding reporter gene FLuc (and later also other reporters) using a blunt-end approach. Each modified 3’-tail sequence (S^1^–S^15^) comprised polyadenine segments of 10 to 45 nucleotides, separated by heteronucleotide spacers of 1 to 6 nucleotides. This design resulted in segmented 3’-tail sequences containing 3 to 12 polyadenine fragments of defined length. The total length of the modified polyA motifs varied from 120 to 206 nucleotides (Fig. 1). In addition, reference molecules representing current polyA tail standards were prepared. Variant R^1^ is a 90 nt polyA tail - the longest unmodified polyA sequence that we could consistently amplify in plasmid DNA without observable shortening. Variant R^2^ contains a modified tail consisting of 30 adenines, a 10-nucleotide spacer (GCATATGACT) followed by 70 adenines, and is used in the Pfizer-BioNTech BNT162b2 mRNA vaccine.(39) Variant R^3^ is a 90 nt polyA tail ending with a pentanucleotide motif (TCTAG) as found in the Moderna mRNA-1273 vaccine.(15) Variants R^4^ and R^5^ consist of 60 adenines, a guanine/cytosine spacer (G/C), followed by another 60 adenines, and have been reported to increase the translational activity of mRNA in cultured cells.(29) Finally, variant R^6^, containing a bulky G-quadruplex structure, was designed as a negative control reference.(40)

**Figure 1.**
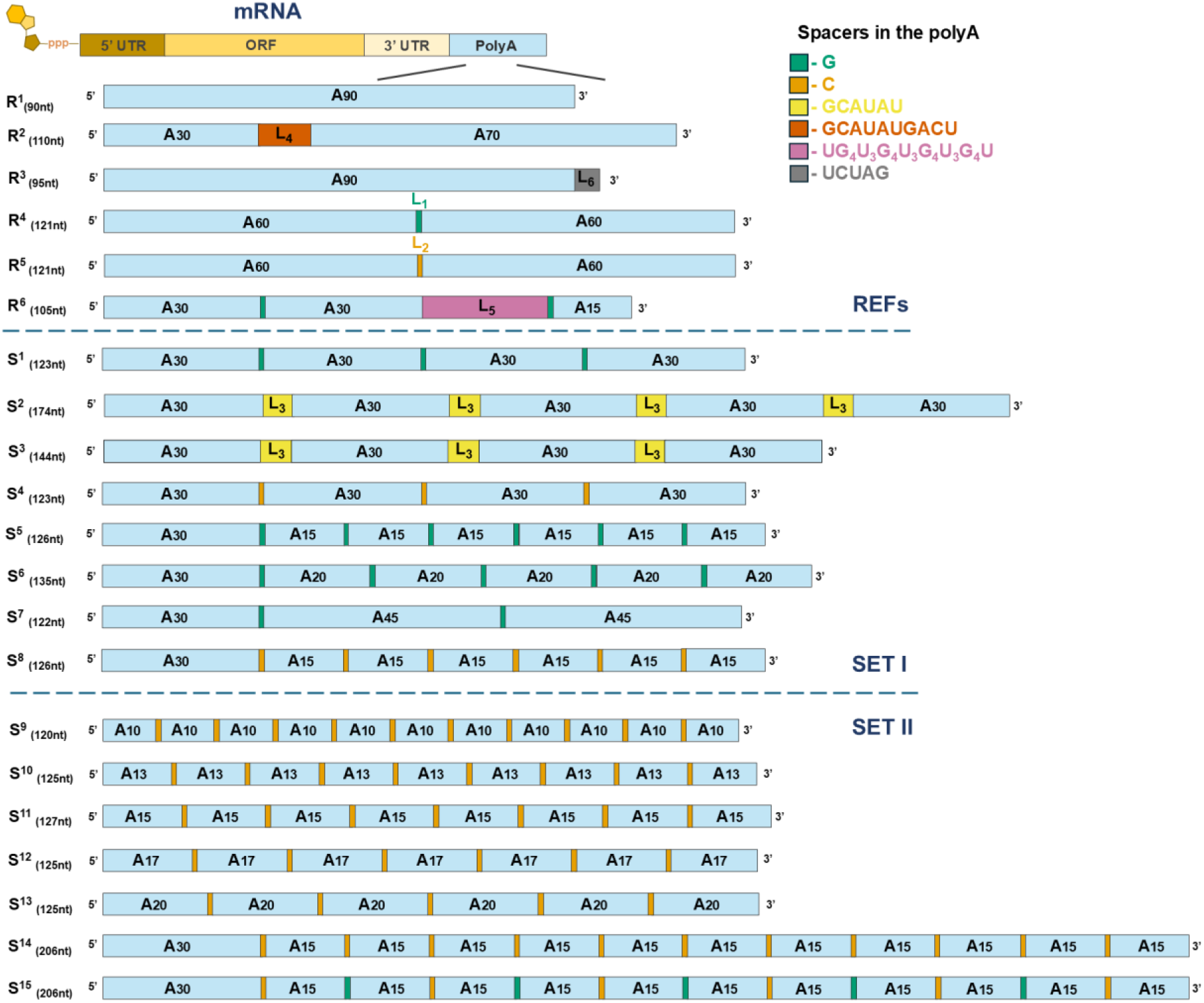
PolyA variants designed and evaluated in this study. R^1^-R^5^ are the reference polyA structures known from previous studies or commercially available mRNA vaccines. R^6^ is an negative control reference that is expected to decrease translational activity. S^1^-S^15^ are the novel modified polyA variants that were designed and evaluated in this study. Labels: adenine (A), guanine (G), cytosine (C), uridine (U) nucleotides. The sign below the label determines the number of nucleotide units in the particular fragment. The evaluated polyA sequences were classified into two sets. Set I includes nucleotide modifications that vary in the type and distribution of spacers within the polyA sequence. Set II, which was developed later, primarily consists of modifications featuring multiple mononucleotide cytosine spacers, that segment the polyA sequence into evenly spaced adenine fragments.

### Stability analysis of plasmid DNA with modified polyA sequences

The stability of the designed RNA sequences in the DNA constructs is crucial, because any alterations to the sequences during amplification of the template can significantly affect the quality and functionality of the resulting mRNA. PolyA tail shortening during plasmid amplification in bacterial systems, which is often caused by DNA polymerase errors, results in constructs with truncated polyA tails.(41,42) The presence of such constructs can decrease the homogeneity of the final mRNA product and can adversely affect its translational efficiency.

Therefore, modifications that exhibit high stability during plasmid DNA propagation are highly desirable. To this end, we conducted a comprehensive analysis of plasmid DNA vectors encoding reporter genes with modified 3’-tail polyA sequences, evaluating their propensity for polyA tail shortening during amplification in two model bacterial systems. The stability of each modified variant S^1^–S^8^ and reference construct R^1^–R^6^, was assessed in two commonly used bacterial systems: *E. coli* Top10 and NEB^®^ Stable Competent (SC) *E. coli* (High Efficiency) using Sanger DNA sequencing of plasmids purified from 30 independently isolated bacterial colonies. The results were expressed as the relative instability, i.e. the percentage of colonies with altered (shortened) sequences compared to the original constructs (Figure 2A). We also included a plasmid containing a polyA tail of 130 adenines in this analysis. However, as expected, it exhibited the highest mutation frequency in tail length and sequence (∼74% and 39% for the TOP10 and SC strains, respectively), which hindered the generation of a stable construct. Therefore, the construct was excluded from further studies. In contrast, the classical polyA tail (R^1^, 90 adenines) and references R^2^ and R^3^ showed greater stability with an instability parameter of 15–17% in the TOP10 strain. References R^4^ and R^5^, which had a single heteronucleotide inserted after 60 adenines, exhibited reduced stability, accounting for 33– 34% of mutated clones in the TOP10. Meanwhile, constructs with the designed modified 3’-tail sequences (i.e. S^1^–S^8^) demonstrated high fidelity and stability comparable to that of R^1^–R^3^ (11–15% and 7–8% for TOP10 and SC strains, respectively). Overall, using NEB^®^ Stable Competent *E. coli* reduced polyA tail loss by ∼2–3-fold during amplification (Fig. 2A). Similar stability of modified sequences was later analogously confirmed for the second set of templates (Fig. S2).

**Figure 2.**
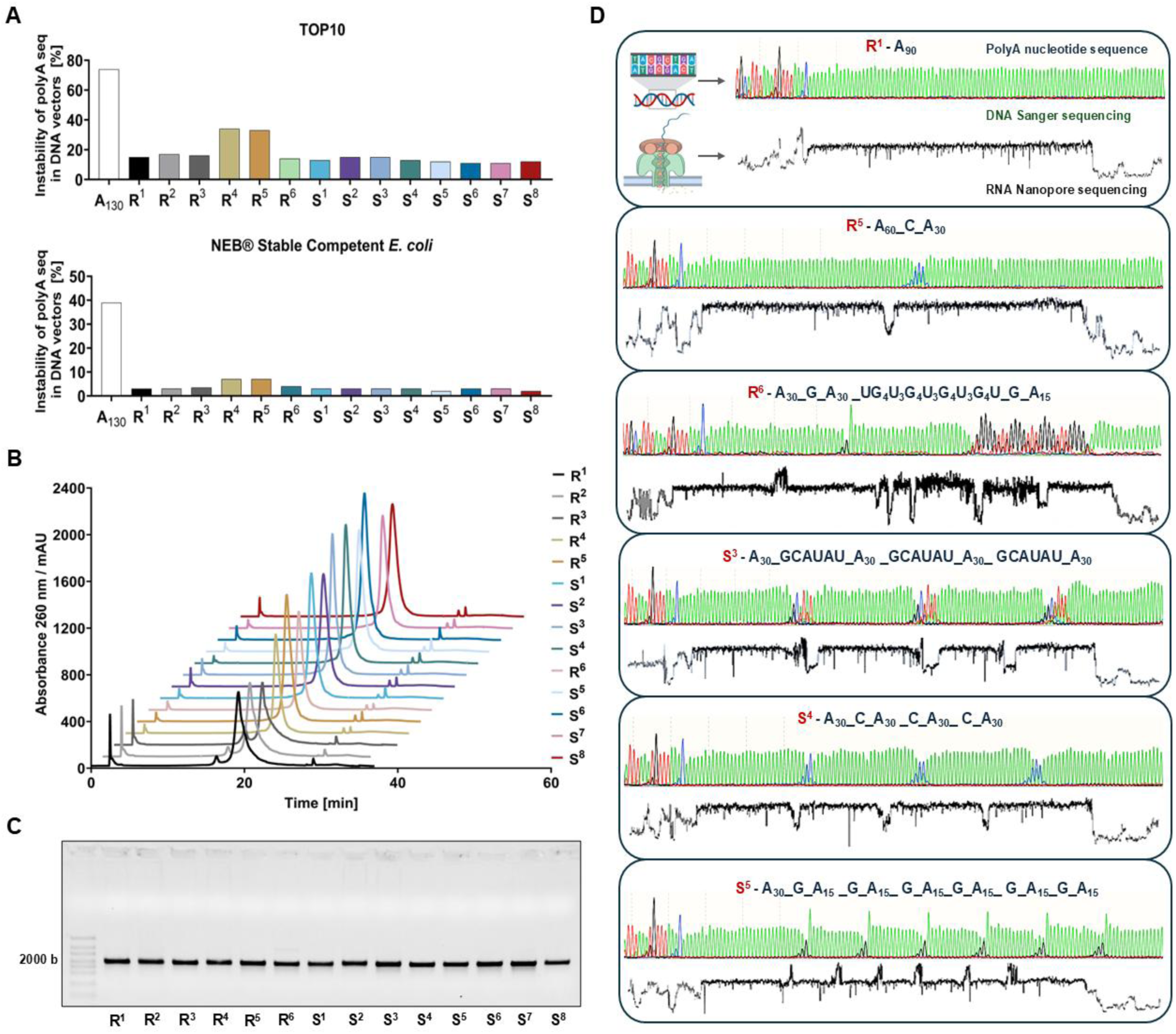
Identity confirmation and stability assessment of modified polyA sequences at the DNA and RNA levels. **A)** Comparison of polyA tail sequence instability levels in plasmid DNA vectors encoding FLuc mRNA with various polyA variants across two bacterial strains: TOP10 and NEB^®^ Stable Competent *E. coli*. Instability levels were expressed as the percentage of clones with altered polyA tail length or sequence relative to the original construct. **B)** Chromatographic profiles of mRNA with modified polyA sequences. Absorbance at 260 nm shows similar elution patterns across polyA constructs, confirming that sequence variations and spacer modifications do not affect mRNA synthesis. **C)** Agarose gel electrophoresis of purified mRNA. **D)** Confirmation of polyA structures at the DNA and RNA level. Comparison of plasmid DNA and IVT RNA sequencing for constructs R^1^, R^6^, S^4^, and S^8^. For clarity, only data for polyA fragments are shown. DNA sequencing was performed by the Sanger method; raw data shows individual nucleotide reads in the polyA region of a plasmid DNA construct encoding Firefly luciferase. Color coding: green-adenine, red-thymine, blue-cytosine, black-guanine. RNA sequencing was performed using the Nanopore DRS method for mRNAs obtained from templates generated by linearization of the plasmid DNA. The results are expressed as changes in current [pA] as a function of sequencing time [s]. Each significant change in the current indicates the occurrence of a heteronucleotide relative to adenine.

### Validation of polyA segmentation patterns at the DNA and RNA levels

The plasmids encoding mRNAs with segmented polyA tails were amplified, purified, and subjected to Sanger DNA sequencing (selected examples are shown in Fig. 2D, all examples are shown in Fig. S1). The analyses confirmed the presence of the designed segmented polyA patterns, with limited heterogeneity. Selected DNA sequences were additionally analysed by direct Nanopore sequencing, confirming the results (Fig. S3). The plasmids were then linearized and used as templates for *in vitro* transcription (IVT) to synthesize capped mRNA encoding firefly luciferase (FLuc). The IVT synthesis yield exceeded 3 mg of crude mRNA per 1 mL of reaction for all tested modifications, remaining consistent with the outputs observed for the reference sequences. The crude mRNA was purified using Poros oligo(dT)_25_ resin, followed by RP-HPLC. The applied purification strategy ensured high purity and homogeneity of the final mRNA preparations, as demonstrated by chromatographic profiles (Fig. 2B) and electrophoretic analysis (Fig. 2C). The polyA sequence modifications did not adversely affect mRNA synthesis efficiency or purification, enabling the production of high-quality mRNA preparations. To verify the accuracy of polyA modifications at the RNA level, direct RNA sequencing (DRS) using Nanopore technology was performed on all mRNA variants with modified 3’-tail sequences. This approach provided a high-resolution analysis of polyA tail sequences with selected examples shown in Fig. 2D and a complete dataset in Fig. S4. Characteristic current shifts induced by heteronucleotide insertions enabled precise identification of spacer types and polyA fragment lengths. Mononucleotide cytosine and guanine spacers exhibited opposite current intensity shifts, while longer spacers produced a distinct intensity pattern. DRS confirmed the presence of designed nucleotide spacers in all modified mRNA variants (S^1^–S^8^) and validated the accuracy of reference sequences. The analysis demonstrated full concordance between mRNA sequences with modified 3’-tail regions and their corresponding DNA templates. Dot-blot analyses for double-stranded RNA (dsRNA) contamination detected no measurable amounts of dsRNA in the modified samples, confirming that structural modifications of the polyA tail did not induce increased dsRNA formation (Fig. S5).

### *In vitro* kinetics and expression of FLuc

mRNA molecules encoding FLuc with modified 3’-tail sequences S^1^–S^8^, along with all reference variants, were transfected into cultured cells to assess their translational activity at various time points. A preliminary analysis of selected polyA-modified mRNAs was conducted to compare translational activities of 1-methyl-pseudouridine-(m^1^Ψ) and uridine-containing (U) mRNAs in HEK293T cells. For this purpose, mRNA for four variants (R^1^, S^1^, S^2^, and S^4^) was synthesized in two versions—containing either canonical uridine (U) or the modified base N1-methylpseudouridine (m¹Ψ). Incorporation of m¹Ψ is known to improve the properties of in vitro–transcribed mRNA by reducing dsRNA formation and consequently innate immune recognition.(43) Consistently, we observed that 3′-modified mRNA variants incorporating m¹Ψ exhibited higher homogeneity after purification and yielded increased FLuc protein expression levels across all polyA tail modifications (Fig. S6). Based on these observations, we selected m¹Ψ-modified mRNA for subsequent experiments. Next, FLuc expression was analysed in four cell lines (A549, HEK293T, HepG2, and JAWS II), representing cancerous and immune cell types from human and murine origins. We found that mRNA translation in these cells was sensitive to polyA tail length (A_30_ versus A_90_; Fig. S7). All m^1^Ψ-mRNA variants from SET I (S^1^– S^8^) and references R^1^–R^6^ were transfected into cells using lipofectamine, and chemiluminescence measurements were performed after 4, 16, 24, 48, and 72 h to obtain the FLuc expression profiles (Fig. 3A–D) and overall protein activities (Fig. 3E–H). The FLuc expression at individual timepoints followed a characteristic pattern for most tested cell lines, with a rapid initial increase in luminescence within the first 4 to 16 hours, peaking at 24 to 48 hours, and subsequently declining in expression levels. However, the magnitude and duration of protein expression varied, depending on both the cell type and the specific polyA modification, whereas cell viability remained unaffected (Fig. S8). Notably, JAWS II cells (murine dendritic cells) displayed a distinct expression profile, with the highest luminescence consistently observed at the earliest time point, 4 hours post-transfection (Fig. 3D). Comparing FLuc activity produced from different mRNA variants revealed that 3’-terminally modified mRNA, particularly variants S^2^, S^4^, S^5^, S^6^, and S^8^, consistently exhibited higher protein outputs than the reference constructs [R^1^ (A_90_), R^2^, and R^3^], regardless of cell type or measurement time point. References R^4^ and R^5^, containing a single heteronucleotide spacer, also outperformed R^1^ and R^2^, whereas R^6^, containing a G-quadruplex structure, had a highly reduced translational activity. Comparison of R^4^ and R^5^ with the modified variants (S^2^, S^4^, S^5^, S^6^, and S^8^) revealed that most of the novel 3’-terminally modified sequences yielded higher luciferase activity. However, at certain time points in the HepG2 and JAWSII cell lines, the expression levels of R^5^ were comparable to those of the novel modified constructs. The advantages of our novel 3’-tail modifications were particularly evident at later post-transfection stages, when modified mRNA variants continued to produce protein. Importantly, the most efficient constructs (S^2^, S^4^, S^5^, S^6^ and S^8^) produced higher protein levels than R^4^ and R^5^. S^4^ and S^8^ exhibited stable protein expression throughout the experiment, indirectly suggesting that these modifications may enhance both translation efficiency and/or mRNA stability. A comparative analysis of total protein production further underscored the advantages of the modified 3’-tail sequences (Fig. 3E–H). In A549 and HEK293T cells, variants S^4^, S^5^, S^6^, and S^8^ produced the highest luciferase activity, with signal intensities 3-to 5-fold greater than those of the reference construct R^1^. In HepG2 cells, S^8^ demonstrated the highest overall activity, exceeding R^1^ by more than 3-fold. While no significant differences in total FLuc activity were observed between R^1^, R^2^, and R^3^, variant R^4^ consistently produced 1.5-to 2-fold higher protein levels than the former references. R^5^ outperformed R^4^, consistently with previous findings (29). In JAWS II cells, S^4^ and S^8^ displayed the highest luminescence levels among all the constructs tested, generating signals that were approximately 2.5 times greater than the A_90_ tail (R^1^). Overall, the modified 3’-tail sequences resulted in a 2-to 5-fold increase in total protein production compared to the non-segmented 90-base long polyA tail (R^1^). Among these, S^4^, S^5^, and S^8^ consistently showed the most beneficial properties, surpassing even the top-performing reference variant, R^5^ (Fig. 3E–H). Statistical analysis confirmed a significant enhancement in translational efficiency for S^4^ and S^8^ compared to R^1^ in HepG2, HEK293T, and JAWS II cells. Additionally, S^8^ demonstrated an advantage in A549 cells, further highlighting its enhanced performance.

**Figure 3.**
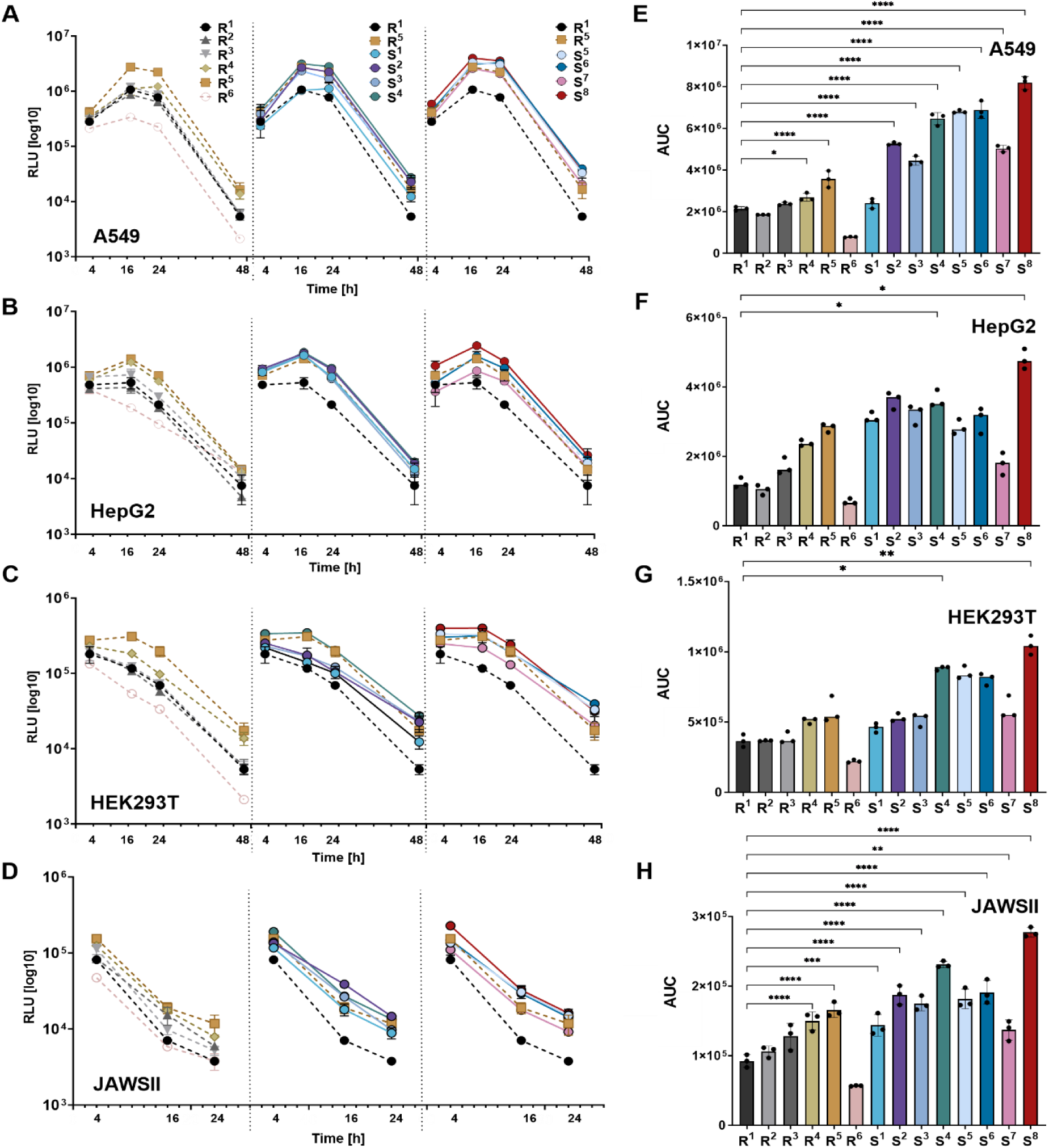
The composition of the polyA tail affects the expression analysis of the encoded Firefly luciferase. **A-D)** Time-dependent production of Firefly luciferase (FLuc) in cultured cells transfected with mRNAs carrying various polyA tails. Graphs show relative activity (chemiluminescence) of FLuc (which is proportional to luciferase protein levels) in cells following transfection with different mRNA variants, including all references (R^1^-R^6^) as well as mRNAs with modified 3’-tail sequences (S^1^-S^8^). Relative activity of Firefly luciferase was measured in four cell lines: A549 (A), HepG2 (B), HEK293T (C), and JAWSII (D) at four time points post-transfection (4 h, 16 h, 24 h, and 48 h). **E-H)** Overall Firefly luciferase production in different cell lines for each mRNA variant relative to reference mRNA (R^1^) represented as the AUC (Area Under the Curve). Graphs show the total FLuc protein from all time points normalized to the same data for R^1^. E) The total FLuc protein expression in the A549 cell line with statistical significance relative to R^1^ (A90). F) The total FLuc protein expression in HEK293T cell line with statistical significance relative to R^1^ (A90). G) The total FLuc protein expression in the HepG2 cell line with statistical significance relative to R^1^ (A90). H) The total FLuc protein expression in the JAWS II cell line with statistical significance relative to R^1^ (A90). Statistical significance was assessed using the Kruskal-Wallis test with Dun’s multiple comparisons test (medians) for HepG2 and HEK293T, (ns,* -p< 0.05,**-p<0.01), and using one-way ANOVA (mean ± SD) for A549 and JAWSII (*p < 0.05, **p < 0.01, ***p < 0.001, ****p < 0.0001).

### *In vitro* stability and protein expression of mKate2_PEST

To elucidate the influence of modified 3′-terminal mRNA sequences on transcript stability and translational efficiency, we employed mRNAs encoding the red fluorescent reporter protein mKate2 fused with a PEST sequence. This well-characterized degradation motif accelerates intracellular protein turnover.(44) The use of this rapidly degraded reporter enabled precise temporal analysis of mRNA translation and stability dynamics. Experiments were conducted in A549 and HEK293T cell lines. All mRNA variants (S^1^–S^8^) and reference constructs (R^1^–R^6^) were prepared and validated analogously as FLuc mRNAs, transfected into cells, and mKate expression was followed by automated fluorescence microscopy at 2-hour intervals from 5 to 77 hours post-transfection. The resulting fluorescence profiles of mKate2_PEST (Fig. 4A) revealed consistent expression kinetics across all constructs, with rapid signal accumulation peaking around 18 hours, followed by a progressive decline and near-complete loss of fluorescence by 50 hours. Compared to A549, HEK293T cells exhibited lower peak fluorescence and a more gradual post-peak decline, indicating potential differences in mRNA transfection rates, stability, or translation dynamics. Mathematical modelling enabled the estimation of mRNA half-lives and translational efficiencies, defined as the initial slope of protein accumulation curves (Fig. 4B, Table S3). The analysis revealed distinct cell line– specific expression dynamics: A549 cells exhibited longer mRNA half-lives (t_0.5_ ∼400 min) and slower protein degradation (t_0.5_ ∼260 min), leading to greater cumulative fluorescence despite lower translational efficiency. Conversely, HEK293T cells displayed faster mRNA turnover (t_0.5_ ∼350 min), shorter protein half-life (t_0.5_ ∼180 min), but higher translational efficiency. Fluorescence quantification (Fig. 4C) in A549 cells revealed that variants S^5^, S^6^, and S^8^ produced 2–3 fold higher total expression (AUC) compared to references R^1^–R^5^, with S^6^ showing the highest mKate2 yield, consistent with its enhanced stability parameters. In HEK293T cells, S^1^ and S^6^ also outperformed R¹ despite overall lower expression levels. Interestingly, R^3^ exhibited fluorescence comparable to top-performing variants, albeit with greater variability. R^6^, which contains a G-quadruplex, showed the lowest expression, consistent with the results obtained for the FLuc reporter. Surprisingly, these results revealed no significant differences in mRNA stability across most modified variants (except R_6_, for which the half-life was significantly shorter), challenging the assumption that the polyA segmentation uniformly stabilizes mRNA. Instead, the increased protein output appears to result from alternative mechanisms, underscoring the complexity of mRNA–protein regulatory relationships.

**Figure 4.**
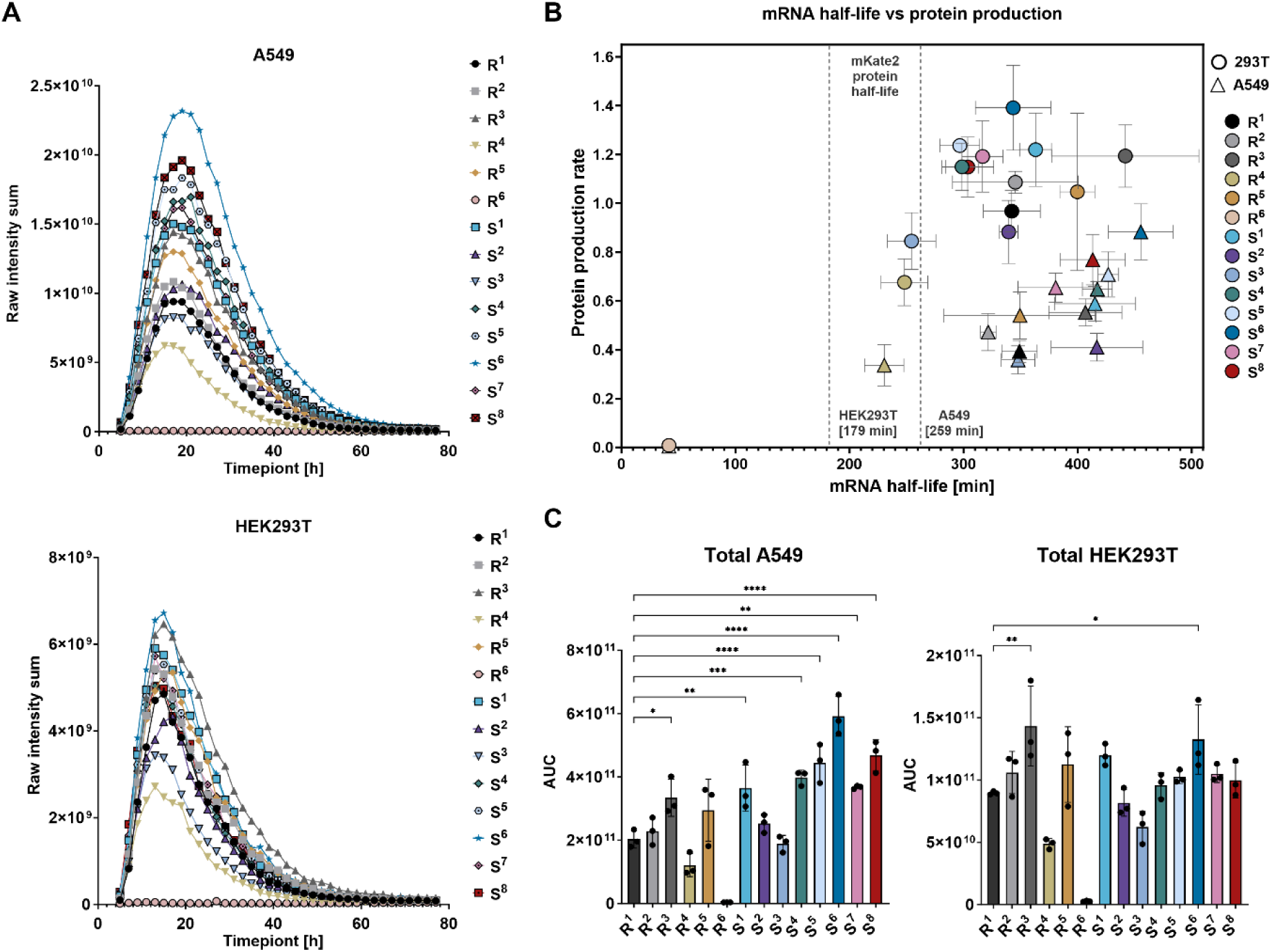
Analysis of mRNAs with polyA tail variants encoding mKate2_PEST: expression kinetics and stability. **A)** Time-dependent expression profiles of mKate2_PEST in A549 and HEK293T cells transfected with mRNAs carrying different modified 3’ terminal sequences. Protein expression levels are presented as the sum of raw fluorescence intensities measured over multiple time points (5–77 h). **B)** Correlation between mRNA half-life and translation efficiency in HEK293T and A549 cells. The graph depicts the half-life of mRNAs encoding mKate2_PEST (X-axis) in relation to the protein synthesis efficiency per mRNA molecule (Y-axis). Each point corresponds to a specific 3’-tail sequence variant, with circles representing HEK293T cells and triangles indicating A549 cells. Dashed lines indicate the half-life of mKate2_PEST in HEK293T (179 min) and A549 (259 min) cells. All values are presented as means ± standard deviation (SD). **C)** Total mKate2_PEST expression levels in A549 and HEK293T cells, represented as the AUC (Area Under the Curve). AUC values were calculated as the sum of the areas under the fluorescence intensity curves, providing a comparative measure of overall protein production for each mRNA variant. Statistical significance was assessed using one-way ANOVA (*p < 0.05, **p < 0.01, ***p < 0.001, ****p < 0.0001).

### Evaluation of polyA modifications with multiple cytosine spacers and extended length in FLuc mRNA

Among the explored polyA patterns, variant S^8^ – comprising A_30_ tract followed by six consecutive CA_15_ tracts – was consistently found among the top-performing ones, regardless of cell type and reporter used. This observation prompted us to further investigate densely modified polyA tails and to design seven additional variants (S^9^–S^15^). Variants S^9^–S^13^ featured a defined and constant number of adenines per motif, separated by single-nucleotide cytosine spacers, but lacked the uninterrupted A_30_ tract present in sequences S^1^–S^8^. In contrast, S^14^ and S^15^ were designed with an extended total polyA length exceeding 200 nucleotides and retained the initial A_30_ motif. In S^14^, cytosine spacers were introduced every 15 adenines, mirroring the structure of S^8^, whereas in S^15^, an alternating pattern of cytosine and guanosine spacers was incorporated every 15 adenines (Fig. 1). The mRNAs encoding FLuc carrying polyA variants S^9^–S^15^, along with previously tested variants R^1^, R^5^, and S^8^ were synthesized, purified, and validated by sequencing as described above (Fig. S1B and Fig S2).

Across the tested sequences, several mRNA variants exhibited either comparable or enhanced expression relative to S^8^, which previously showed one of the highest expression levels. Variants R^1^–R^6^ and S^9^–S^12^ displayed comparable or lower FLuc activity than S^8^, whereas S^13^–S^15^ demonstrated markedly elevated and prolonged expression. These differences appear to correlate with both the presence and spacing of cytosine spacers as well as the overall length of the polyA stretch. Notably, even densely modified constructs such as S^9^ and S^10^ – featuring frequent cytosine interruptions within the polyA tail – remained translationally active, suggesting that extensive polyA modifications do not inherently block ribosome recruitment or otherwise impair protein synthesis. We believe that this observation challenges the prevailing view of polyA modification limits and highlights new possibilities for further exploration of genetic and chemical alterations of polyA tail. Most interestingly, variants S^13^, S^14^, and S^15^ maintained elevated expression for extended durations, indicating that their specific sequence architecture may enhance translational efficiency and/or reduce mRNA degradation, thereby sustaining protein output. This effect was particularly pronounced in HEK293T and A549 cells, where detectable luciferase activity persisted even at later time points (48–72 hours post-transfection). Time-course analysis of FLuc expression further confirmed that constructs incorporating cytosine spacers at defined intervals—most notably S^13^, with a cytosine inserted every 20 adenosines—and constructs with extended polyA tails (S^14^ and S^15^), supported significantly higher and more durable protein production compared to both the canonical polyA tail (R^1^) and the single-cytosine spacer variant (R^5^) (Fig. 5A–D).

**Figure 5.**
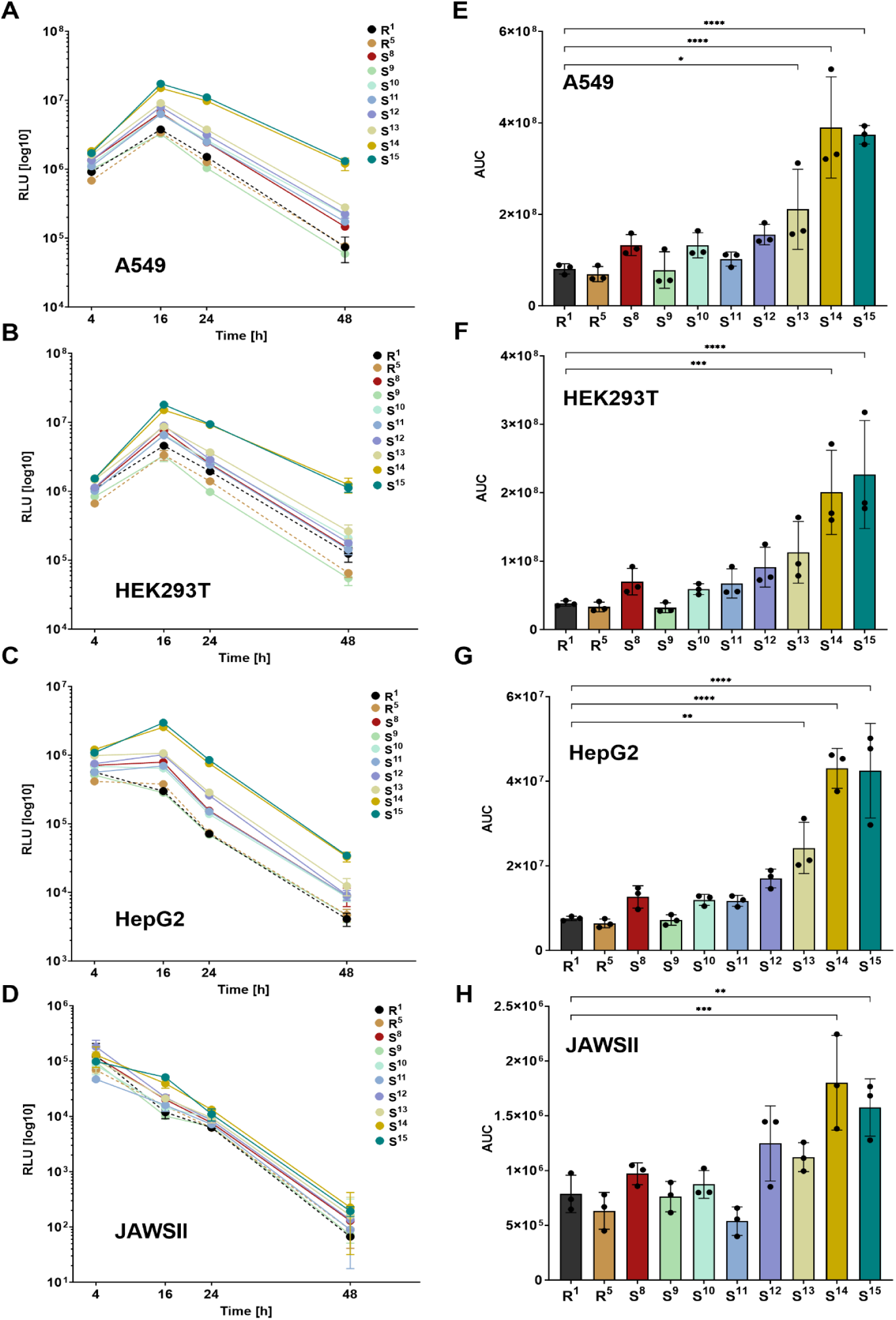
Overall FLuc production in different cell lines for the polyA modifications with multiple cytosine spacers and extended length. **A–D)** Time-course analysis of Firefly luciferase (FLuc) expression in cell lines: A549 (A), HEK 293T (B), HepG2 (C), and JAWSII (D). The graphs illustrate FLuc protein expression for polyA tail modifications (variants: S^9^, S^10^, S^11^, S^12^, S^13^, S^14^, and S^15^) compared to mRNAs carrying the canonical polyA sequences R^1^ and R^5^, as well as the best variant from SET I - S^8^. **E–H)** Area under the curve (AUC) analysis of FLuc protein production over time for each polyA tail modification in A549 (E), HEK 293T (F), HepG2 (G), and JAWSII (H) cell lines. AUC values were calculated by summing the areas under the curves (A–D) at each time point. All values are presented as mean ± standard deviation (SD). Statistical significance was assessed using one-way ANOVA (*p < 0.05, **p < 0.01, ***p < 0.001, ****p < 0.0001).

Quantifying the overall protein output over time further supports these observations (Fig. 5 E–H). In all four cell lines, S^13^, S^14^, and S^15^ consistently exhibited the highest cumulative FLuc expression, confirming their superior translational performance. Importantly, S^14^ and S^15^ displayed a 4-to 6-fold increase in total protein production compared to the reference sequence R^1^, with S^14^ emerging as the top-performing variant across nearly all tested cell lines. Specifically, S^14^ exhibited an approximately 6-fold increase in cumulative FLuc expression compared to R^1^ in A549 cells, as well as a similar enhancement in HEK 293T, HepG2, and JAWSII cells. S^13^ resulted in a 2-to 3-fold increase in FLuc production relative to R^1^. While S^13^ outperformed the other shorter segmented polyA variants (S^9^–S^12^), its expression levels remained below those of the extended polyA modifications (S^14^ and S^15^). This suggests that both the segmentation pattern and total polyA length contribute to modulating translational efficiency. The reference sequences, R^1^ (canonical polyA) and R^5^ (single-cytosine spacer modification), exhibited the lowest AUC values across all tested conditions, reinforcing the notion that structural modifications within the polyA tail can enhance the function of IVT mRNA. Taken together, these findings confirm that optimized polyA tail segmentation, particularly with periodic cytosine spacers at controlled intervals and extended polyA length, enhances mRNA stability and translational efficiency beyond what is achievable with canonical homopolymeric template-encoded polyA tails. The robust and sustained expression of S^13^–S^15^ across multiple cell types indicates that this approach is broadly applicable and not restricted to a specific cellular context.

The superior performance of S^8^ and S^13^–S^15^ suggests that precise tuning of polyA structure can significantly impact the effectiveness of mRNA therapeutics, vaccines, and gene expression systems, offering new possibilities for enhancing the stability and efficiency of synthetic mRNAs in both research and clinical applications. Hence, to further validate the potential applicability of these modified polyA tails, we tested them in more complex biological models.

### *In vivo* evaluation of 3’ mRNAs with segmented polyA tails

Encouraged by the *in vitro* findings, we aimed to determine whether the observed improvements in mRNA translational activity can be observed *in vivo*. To address this, we selected highly efficient segmentation patterns and tested them in a mouse model, which provides a physiologically relevant system for assessing mRNA stability, biodistribution, and translational potential. For the *in vivo* studies, we utilized two reporter genes, Firefly luciferase (FLuc) and human erythropoietin (hEPO), which enabled quantitative protein expression analysis through bioluminescence imaging (FLuc) and ELISA-based detection (hEPO). Cell-based studies have shown that polyA modifications incorporating cytosine monospacers typically exhibit the highest translational potential. Therefore, we first selected variants S^4^, S^8^, and S^13^, each defined by a distinct cytosine distribution within the polyA tail for *in vivo* evaluation. Additionally, we included the classic A_90_ polyA tail - R^1^ and the most efficient reference - R^5^. All mRNA variants were synthesized as previously described and formulated into lipid nanoparticles (LNPs) using the SM-102-based lipid mix. Dynamic light scattering (DLS) analysis confirmed consistent nanoparticle sizes across the tested modifications, with an average LNP diameter of ∼76 nm for FLuc variants and ∼64 nm for hEPO (Fig. 6A, Fig. S9A). Furthermore, the encapsulation efficiency, ranging from 84% to 92% and PDI ranging from 0.01 to 0.06, indicated the high quality of the formulated LNPs. In the FLuc study, each experimental group consisted of four mice, with mRNA LNPs carrying R^1^, R^5^, S^4^, and S^8^ modifications administered intravenously. *In vivo* bioluminescence imaging was conducted at 4, 8, 12, and 24 h post-administration to monitor protein expression over time (Fig. S10A). The overall expression kinetics were consistent across all tested mRNAs, with the highest bioluminescence signals observed at 4 hours post-administration, followed by a gradual decline over time (Fig. 6B). R^1^ exhibited a slightly slower rate of signal decline after 8 hours; however, this did not substantially alter its overall kinetic profile. Despite consistently displaying the lowest bioluminescence intensities among all tested variants, R^1^ still generated a relatively strong signal, confirming its functional activity *in vivo*. The R^5^ variant followed a similar kinetic trajectory to R^1^, but demonstrated higher bioluminescence intensities at an early 4-hour time point, suggesting a moderate improvement in translational activity. The S^4^ variant exhibited a very similar kinetic profile to R^5^, with comparable bioluminescence intensities across all measured time points. This suggests that the introduction of cytosine spacers every 30 adenines in S^4^ provides a modest translational advantage over the classic polyA tail (R^1^), but does not significantly enhance mRNA translatability beyond that of R^5^. S^8^ exhibited the highest and most sustained bioluminescence signal (Fig. 6D). At 4 hours post-administration, S^8^ expression was markedly higher compared to the other variants, indicating superior translational activity (Fig. 6C). However, despite its higher protein output, the rate of signal decline for S^8^ closely mirrored that of the other variants. This observation suggests that while S^8^ enhances translation efficiency, it most likely has a limited impact on stability *in vivo*.

**Figure 6.**
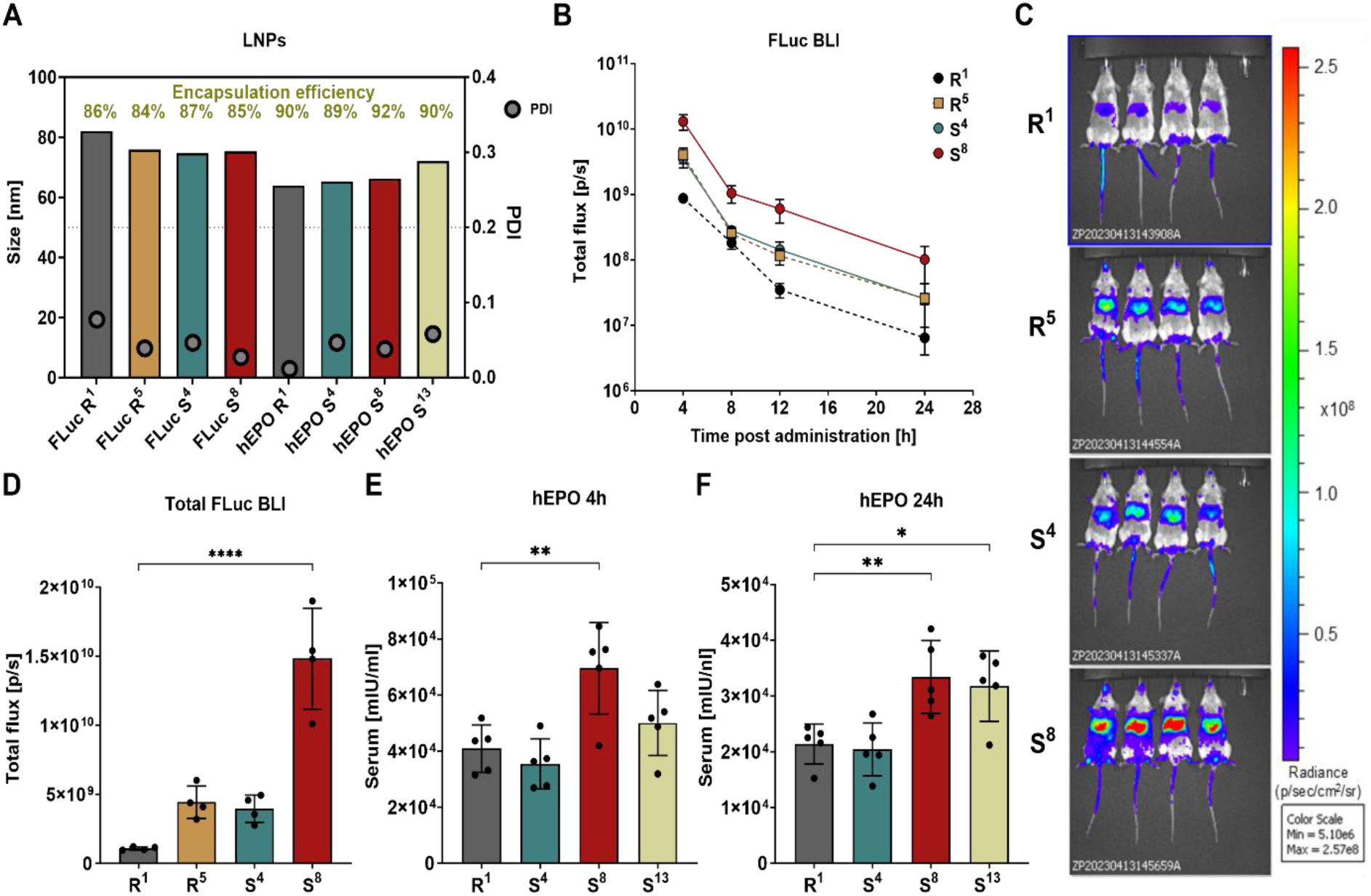
Segmented polyA variant S^8^ [A30(CA15)6] enhances the *in vivo* production of FLuc and hEPO from modified mRNA in a murine model. **A)** Quality control of lipid nanoparticles (LNPs) used in in vivo studies. Average LNP size (Z-average) for all FLuc-and hEPO-encoding mRNAs was determined by dynamic light scattering (DLS). mRNA encapsulation efficiency in SM-102 lipid formulations is shown as the percentage of total RNA encapsulated. Polydispersity index (PDI) values reflect LNP homogeneity; values <0.2 indicate a uniform population. **B)** Quantification of bioluminescence over time (4, 8, 12, and 24 h) following intravenous (i.v.) administration of FLuc-encoding mRNAs (10 µg/mouse) formulated in SM-102 LNPs. n = 4 mice per group. **C)** In vivo bioluminescence imaging at 4 hours post mRNA administration. **D)** Total bioluminescence signals calculated as the sum of bioluminescence measured at 4, 8, 12, and 24 h. All values are presented as mean ± standard deviation (SD). **E, F)** Quantification of serum hEPO concentrations at 4 h (**E**) and 24 h (**F**) post i.v. administration of hEPO mRNA (1 µg/mouse) formulated in SM-102 LNPs. All values are presented as mean ± standard deviation (SD), n = 4 mice per group. Statistical significance was assessed using one-way ANOVA (*p < 0.05, **p < 0.01, ***p < 0.001, ****p < 0.0001).

To further validate these findings with an alternative reporter gene, we generated mRNA molecules encoding human erythropoietin (hEPO) incorporating specific modified polyA sequences. For the hEPO experiment, we chose non-segmented 90-base long polyA tail (R^1^), the previously studied S^4^ and S^8^ variants, as well as S^13^, which demonstrated the highest *in vitro* protein expression among the set II modifications of similar length to S^4^ and S^8^. The mRNA preparations were synthesized and formulated into LNPs following the previously described methodology, ensuring high purity and homogeneity, as confirmed by quality control assessment (Fig. 6A, Fig. S9B). The hEPO-LNPs, incorporating all modifications, were intravenously administered to mice, with each experimental group consisting of five animals. Blood samples were collected at 4 and 24 h post-administration, and serum was collected for quantification of hEPO concentrations using ELISA. (Fig. S10B) The analysis revealed the significant impact of modified 3’-tail sequences on mRNA hEPO translation efficiency and protein expression *in vivo*. At 4 h post-administration (Fig. 6E), the highest hEPO concentrations were observed in the group treated with S^8^-modified mRNA, significantly exceeding those in the R^1^ group and indicating enhanced translational efficiency. In case of S^13^, hEPO concentrations were also higher than those observed with R^1^, although the difference was less pronounced than with S^8^, suggesting a moderate improvement in translation. Conversely, S^4^ did not exhibit a clear difference in protein levels relative to R^1^. At 24 h post-administration (Fig. 6F), the S^8^ modification continued to confer a significant advantage in hEPO production, confirming its ability to support prolonged protein expression. Notably, at this time point, S^13^ also demonstrated a statistically significant increase in hEPO concentrations compared to R^1^, indicating that its effect on translation efficiency becomes more evident over time.

Finally, we compared the *in vivo* translational activity of S^8^ mRNA with its elongated variants – S^14^ and S^15^ – encoding FLuc. Quantitative analysis of bioluminescence (Fig. 7A) revealed that all three modified constructs S^8^, S^14^, and S^15^ exhibited significantly enhanced and sustained bioluminescence signals compared to non-segmented 90-base long polyA tail (R^1^). While the overall expression kinetics were similar across variants, with peak expression observed at 4 hours post-administration, total luminescence levels remained higher for modified constructs throughout all measured time points (4–24 h). These results confirm that the tested polyA segmentation patterns substantially improve translational output *in vivo*. A statistical comparison of total bioluminescence (Fig. 7B) revealed that S^14^ yielded the highest cumulative FLuc signal, reaching levels 6-to 8-fold greater than those of R^1^, with S^8^ and S^15^ also demonstrating increases in protein expression. Detailed kinetic profiles for individual mice (Fig. 7C-E) further supported enhanced translational activity. While S^8^ and S^15^ exhibited some inter-animal variability in signal intensity, S^14^ showed particularly stable and consistently high expression across all time points, highlighting its superior translational stability *in vivo*. Representative bioluminescence imaging at 4 hours post-injection (Fig. 7F) visually confirmed these findings: mice inoculated with LNP-formulated S^14^ mRNA displayed markedly stronger luminescence, with clear localization of the signal in the liver region.

**Figure 7.**
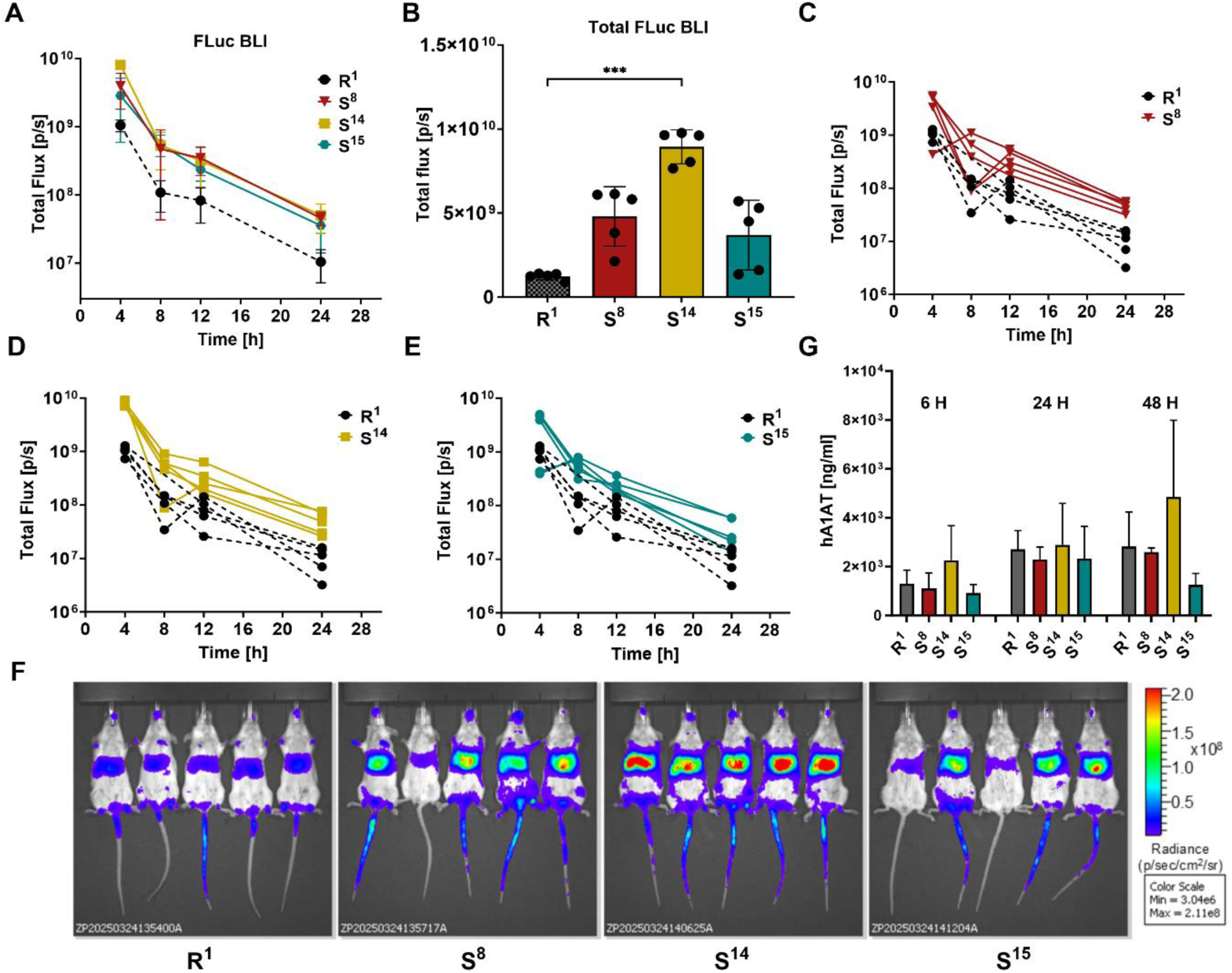
Extended segmented polyA tail (S^14^) further enhances *in vivo* protein production from synthetic mRNA in mice. **A)** Quantification of bioluminescence over time (4, 8, 12, and 24 h) following intravenous (i.v.) administration of FLuc-encoding mRNAs (10 µg/mouse) formulated in SM-102 LNPs. **B)** Total bioluminescence signals calculated as the sum of bioluminescences measured at 4, 8, 12, and 24 h. All values are presented as mean ± standard deviation (SD). **C–E)** Individual bioluminescence profiles of mice inoculated with FLuc mRNA variants. **F)** Representative bioluminescence images taken 4 hours after mRNA administration. **G)** Concentrations of human alpha-1 antitrypsin (hA1AT) in the sera of mice inoculated i.v. with 1 µg of mRNA encoding hA1AT, formulated in SM-102 LNPs. All data are presented as mean ± standard deviation (SD), with *n* = 5 mice per group. Statistical significance was assessed using one-way ANOVA (***p < 0.001).

To further assess the functional relevance of polyA modifications, we extended the analysis to a therapeutic context by testing mRNA encoding human alpha-1 antitrypsin (hA1AT) formulated with selected polyA variants (R^1^, S^8^, S^14^, and S^15^). Serum hA1AT concentrations were measured at 6, 24, and 48 hours post mRNA administration (Fig. 7G). Although inter-individual variability resulted in large standard deviations, S^14^ showed slightly higher protein concentration at 6 and 48 hours after mRNA administration.

## CONCLUSIONS

Structural modifications to the polyA tail can greatly improve the biological effectiveness of synthetic mRNA, making them a key factor in optimizing mRNA-based therapeutics. In this study, we systematically investigated the impact of polyA tail modifications on mRNA translation efficiency and protein expression in different biological contexts. We prepared 45 different genetic constructs, each varying in polyA tail sequence, for four reporter proteins: FLuc, mKate2_PEST, hEPO, and hA1AT. These constructs were designed to examine the effect of specific structural alterations within the polyA tail on mRNA stability, translational efficiency, and overall protein output. We tested both novel and previously described polyA designs, enabling us to make a comprehensive comparison of their functional properties in cell cultures. The therapeutic potential of the most promising modifications was then evaluated using an *in vivo* mouse model.

The first key finding was that all novel modification patterns increased the stability of the corresponding plasmid constructs at the DNA level, compared with polyA sequences of the same length. This feature is a prerequisite for any practical application. Notably, variants S^14^ and S^15^, which comprise tails of more than 200 nt in total, are, to our knowledge, the longest polyA tails incorporated into mRNA by the circular plasmid template-encoded approach. We believe that this approach can be used to create even longer modified tails, getting closer to the limits of enzymatically polyadenylated IVT mRNAs, which can have polyA tails varying in length from up to 500 bases.(45) Our approach also allows us to address the limitations of linear plasmid-based systems such as pJAZZ, which is a powerful tool for cloning repetitive or AT-rich DNA sequences,(46) but requires specialized host strains, yields lower transformation efficiency and relatively low copy number, and requires a large vector backbone size.

Based on protein expression studies, we identified the most promising polyA segmentation patterns. These exhibited significantly higher and more stable protein expression than the classic polyA tail, which is composed exclusively of adenine residues. Of the many tested modifications and variants, S^8^ [A_30_(CA_15_)_6_], S^13^ [A_20_(CA_20_)_5_], and S^14^ [A_30_(CA_15_)_11_] emerged as particularly promising, demonstrating high and stable protein expression *in vitro* and *in vivo*. However, it is worth noting that even modified tails with the most densely packed cytosine spacers [e.g., S^9^ [A_10_(CA_10_)_11_] retained good translational activity compared to R^1^. This indicates that a broader range of modification patterns than previously anticipated can be compatible with polyA function. However, not all modifications resulted in improved performance. For instance, introduction of G-quadruplex structures completely abolished translation (R^6^) and significantly shortened mRNA half-life, confirming that certain sequences in polyA can negatively impact mRNA properties. Contrary to our initial assumption, the data show no clear correlation between modification patterns and mRNA stability. Slight differences in mRNA expression profiles *in vivo*, suggesting higher durability, were observed for the mRNAs with the most extended tails (S^14^ in particular). Comparison of sequences containing 30 consecutive adenines located at the 5’ end of the polyA tail with their segmented counterpart of similar length (e.g. S^8^ versus S^11^) revealed only minor differences in biological activity. This region might be crucial for the efficient recruitment of PABPC1, which requires approximately 27 uninterrupted adenines for effective binding to the polyA tail (35,47). Our data indicate that appropriately designed heteronucleotide modifications of this region may still support the binding. Overall, these results indicate that multiple polyA tail modification patterns are well tolerated by the translational machinery, suggesting that the limits of mRNA tail modification can be extended further – opening new avenues for the design and functionalization of therapeutic mRNAs.

In summary, this study provides new insights into how modifications to the 3’-terminal polyA sequence can enhance the translational efficiency of mRNA molecules. Our findings highlight the potential of polyA tail engineering as a powerful tool for optimizing mRNA-based therapeutics. By introducing specific sequence modifications, it is possible to increase translational activity while simultaneously overcoming the length limitations associated with traditional homopolymeric polyA sequences. The identified extended segmentation patterns (S^14^ and S^15^) have strong therapeutic potential, enabling the sustained expression of therapeutic proteins *in vivo*, which marks a significant advancement in mRNA-based technologies. Further studies are necessary to test if the segmentation approach can produce even longer and more translationally active mRNAs and dissect the underlying mechanism. Notably, the developed polyA fragmentation patterns, such as S^14^, could play a pivotal role in future therapeutic strategies, offering more effective tools for treating a wide range of diseases through gene and molecular therapies.

## DATA AVAILABILITY

Raw sequencing data (fast5 files from Direct RNA Sequencing, nanopore) from RNA reporters are available at Zenodo, doi: 10.5281/zenodo.17376117.

Raw sequencing data (pod5 files) for R^1^, S^14^, and S^15^ plasmids are available at the European Nucleotide Archive (ENA, https://www.ebi.ac.uk/ena/), project accession ERP182322.

Imaging data, together with the processing script, were deposited at the BioImage Archive (EBI, https://www.ebi.ac.uk/bioimage-archive/) under accession: S-BIAD2347

All other raw data are available from the corresponding author upon reasonable request.

## SUPPLEMENTARY DATA

Supplementary Data are available at

## AUTHOR CONTRIBUTIONS

J.K. and J.J. Funding Acquisition. J.K., J.J., and T.S.: Conceptualization. T.S.: Methodology, Validation, Investigation, Visualization, Resources. J.K., and T.S.: Writing—Original Draft. K.C. and Z.P.: Investigation, Validation. D.N.: In vivo data analysis. M.R.B., P.S.K., K.A., W.A., S.M.: Investigation, Visualization, Resources. M.S.G.: Investigation, Resources., M.B., S.Ch.: Resources. D.N., J.G, S.M., A.D., J.J. J.K..: Supervision. All authors: Writing – Review & Editing. All authors have approved the final version of the paper.

---

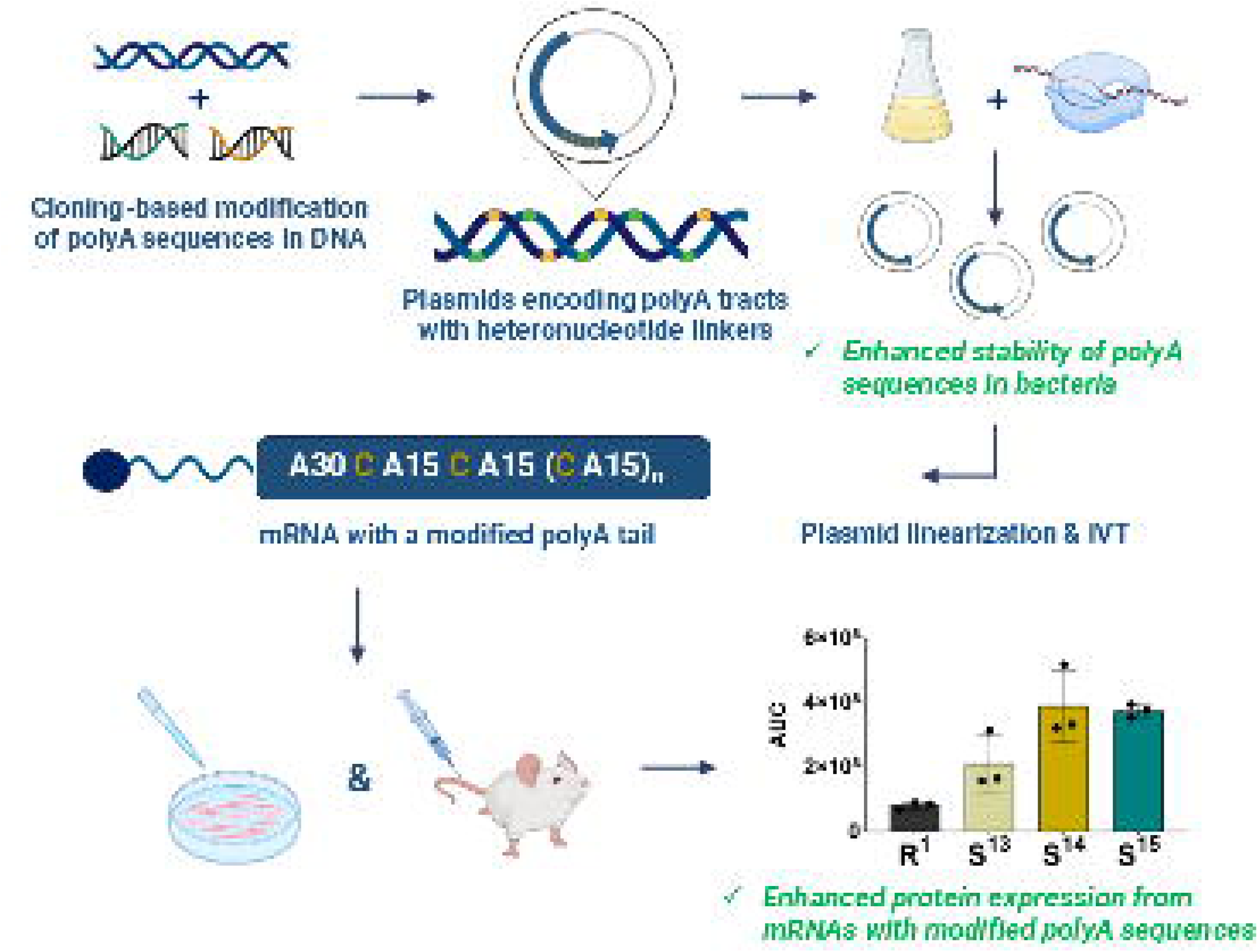

