## Supplementary data for "PolyA tail segmentation improves the stability of the template DNA and increases the translatability of *in vitro* transcribed mRNA"

**SUPPORTING INFORMATION**

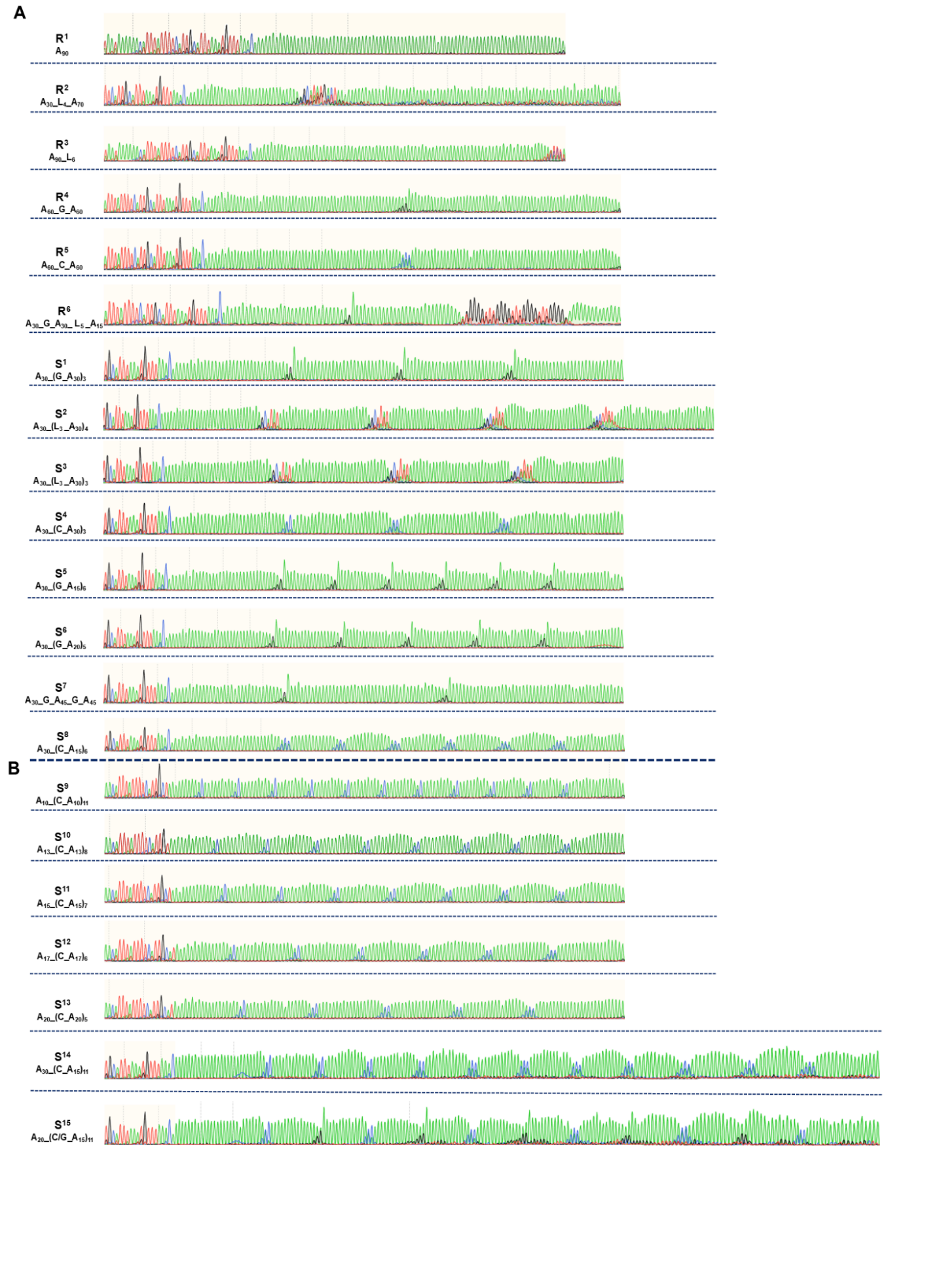

### **Figure S1 Sanger sequencing of modified polyA tail sequences from plasmid DNA encoding FLuc.** The sequencing reads depict the profiles of analysed polyA tail variants, which include nucleotide modifications. Variants R^1^–R^6^ represent reference polyA tail sequences, while S^1^–S^15^ contain various insertions and modifications. Below the modification names, the corresponding summary formulas of the modifications are provided. The coloured peaks correspond to nucleotide bases: adenine (green), cytosine (blue), guanine (black), and thymine (red).

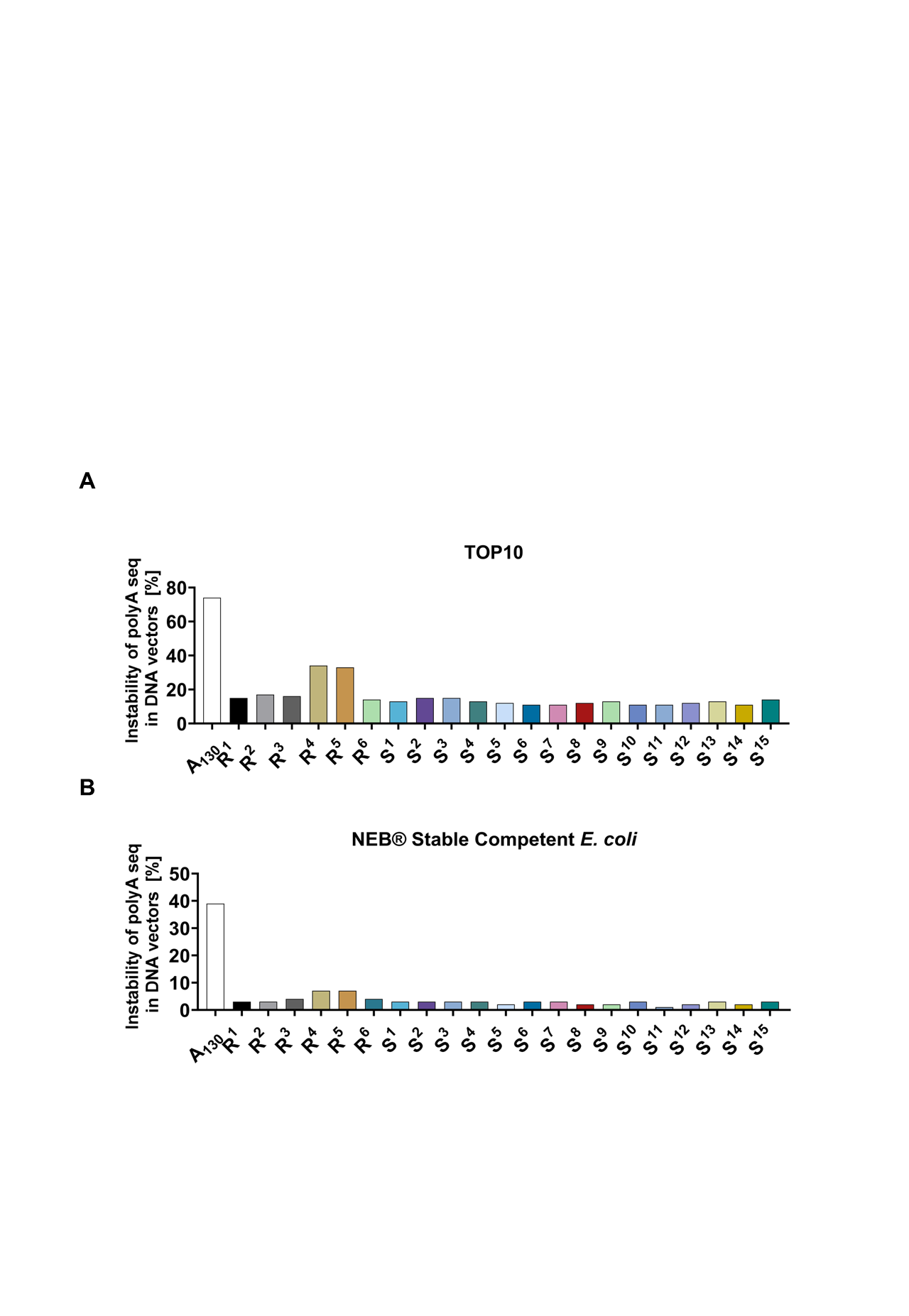

### **Figure S2 PolyA tail sequence instability in plasmid DNA vectors across bacterial strains.** Instability levels of FLuc mRNA polyA variants (R^1^-R^6^, S^1^-S^15^) in TOP10 and NEB® Stable Competent *E. coli* are shown as the percentage of clones with altered polyA tail length or sequence relative to the original construct.

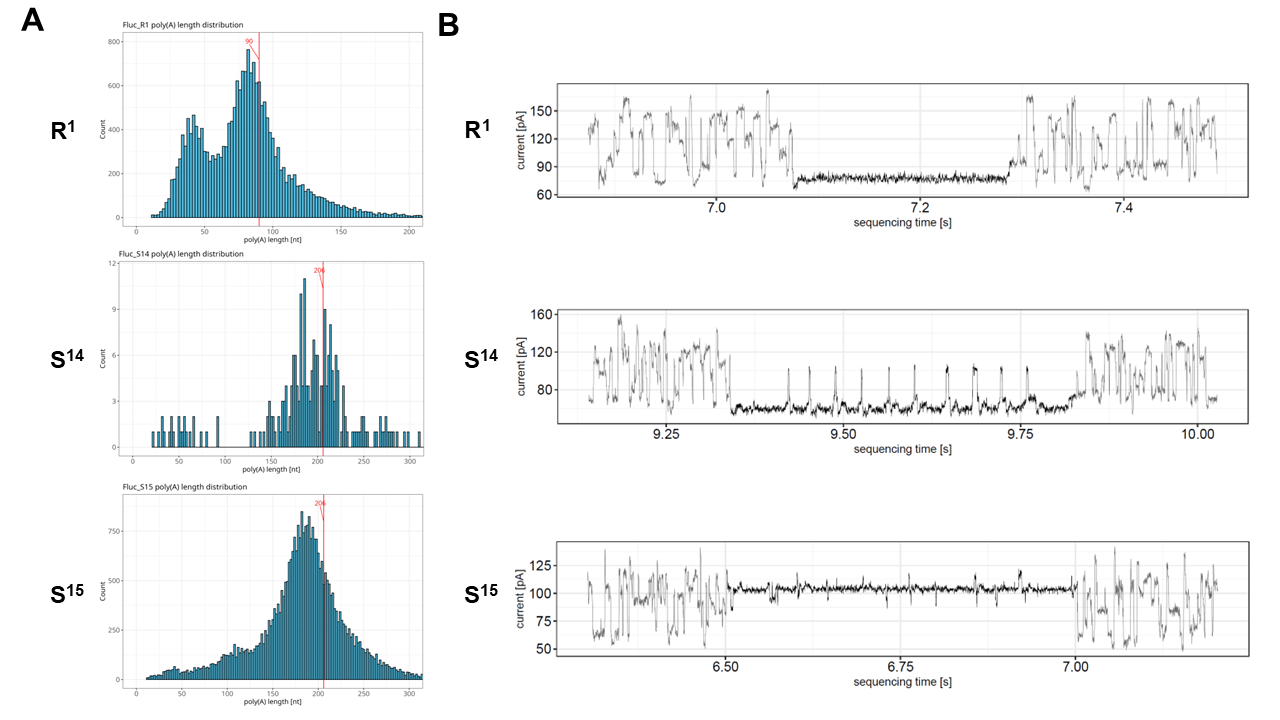

### **Figure S3** **Sequencing reads of DNA Nanopore for selected DNA variants containing modified 3'-tail sequences.** A) Histograms of polyA tail length determined by DNA Nanopore sequencing for selected DNA variants (R^1^, S^14^ and S^15^). B) Changes in current intensity over time for different polyA tail variants with incorporated heteronucleotides. The stepwise changes in intensity reflect the presence of heteronucleotides in the polyA tail structure, which enabled the precise identification of linker types and lengths in the individual DNA variants.

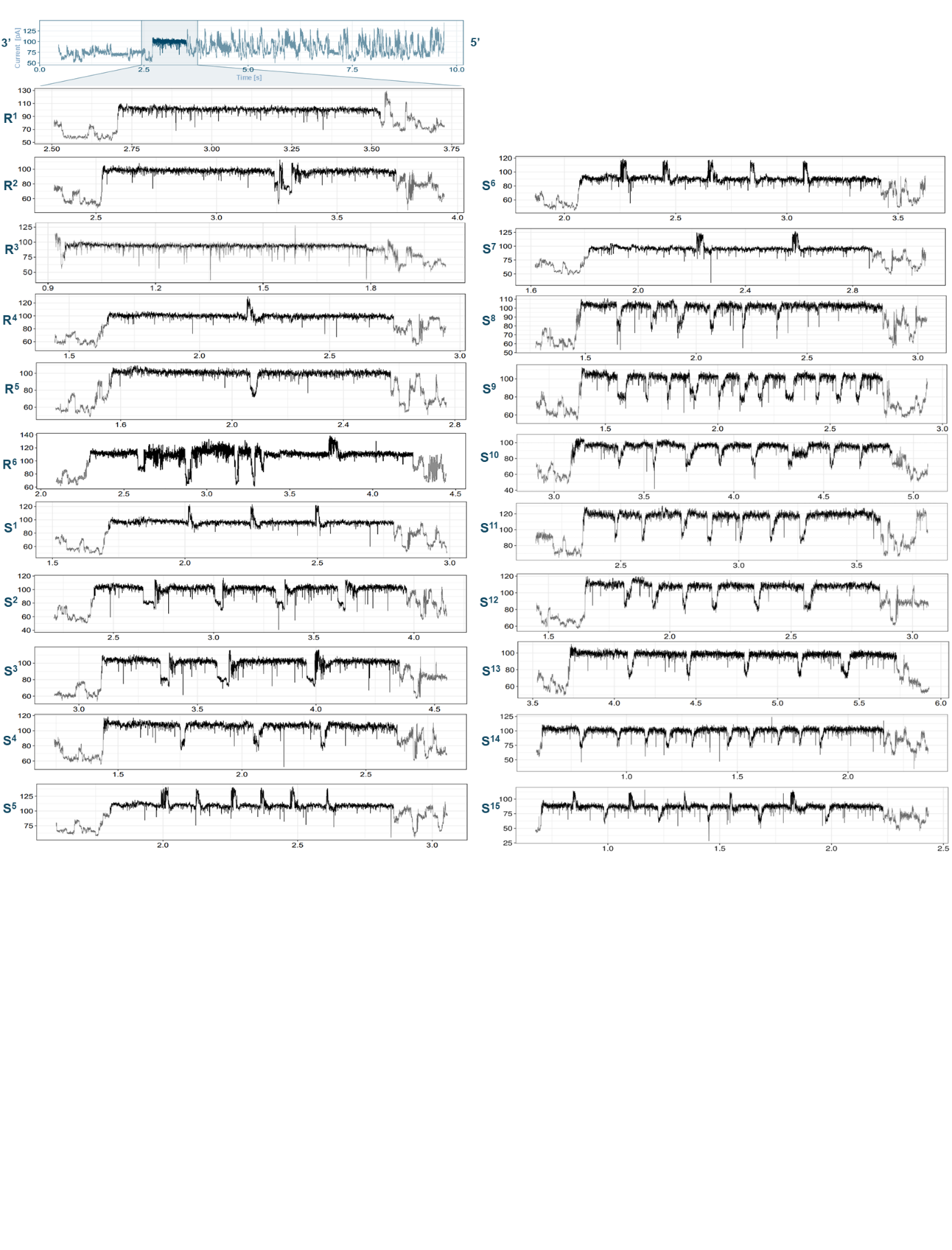

### **Figure S4** **Sequencing reads of DRS for selected mRNA variants containing modified 3'-tail sequences.** Each graph shows changes in current intensity over time for different polyA tail variants with incorporated heteronucleotides. The stepwise changes in intensity reflect the presence of heteronucleotides in the polyA tail structure, which enabled the precise identification of linker types and lengths in the individual mRNA variants.

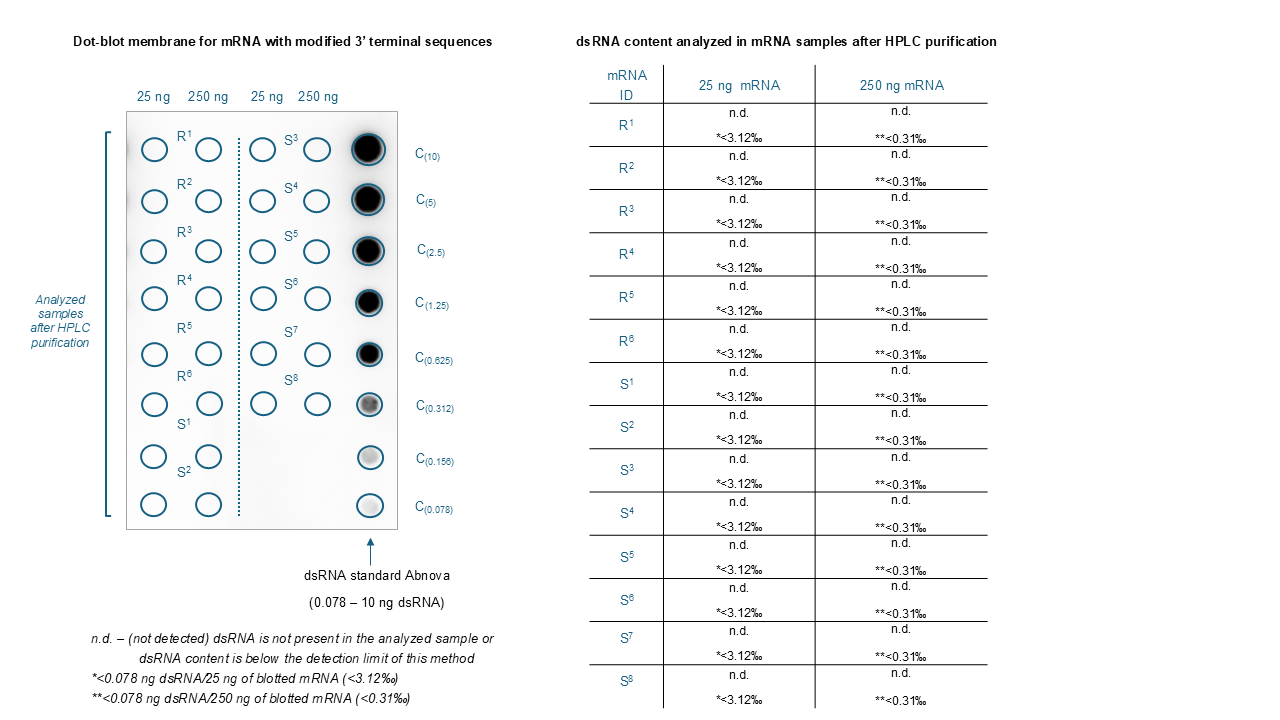

### **Figure S5** **Dot-blot assay results for dsRNA detection in mRNA samples with modified 3'-terminal sequences.** Reference variants are labeled as R^1^–R^6^, while modified variants are designated as S^1^–S^8^. dsRNA levels were determined based on the standard curve range for dsRNA (0.078–10 ng dsRNA). The table provides quantitative results for samples containing 25 ng and 250 ng of mRNA, detailing detection limits (<0.078 ng dsRNA) and presenting results as a percentage of the total mRNA mass.

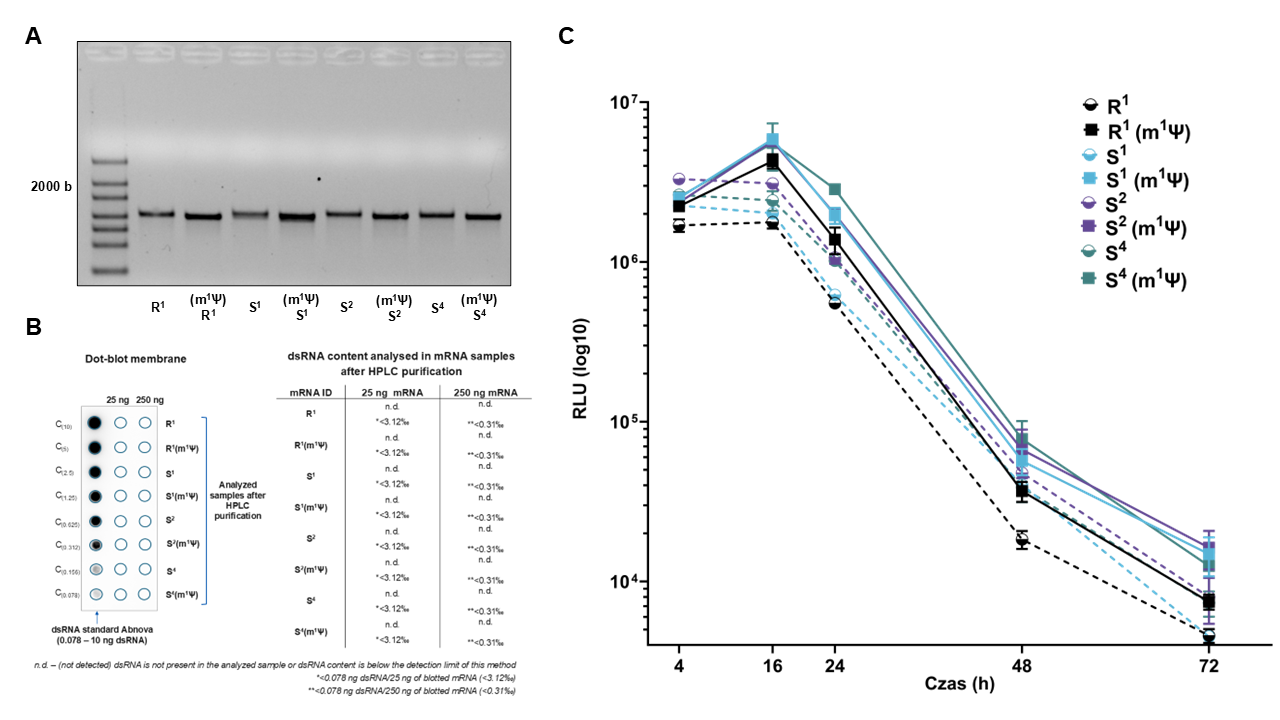

### **Figure S6** **Analysis of mRNA purity and protein expression for variants containing 1-methyl-pseudouridine (m1Ѱ) and uridine (U).** A) Electropherogram of purified mRNA samples, demonstrating high homogeneity and successful incorporation of m1Ѱ. The observed shift in RNA bands for m1Ѱ-modified samples, compared to their U-containing counterparts, confirms the presence of the modification. B) Dot-blot dsRNA analysis of mRNA samples after HPLC purification, showed no detectable levels of double-stranded RNA ( C) FLuc protein expression kinetics in HEK293T cells, measured in relative light units (RLU) at defined time points (4 h, 16 h, 24 h, 48 h, and 72 h).

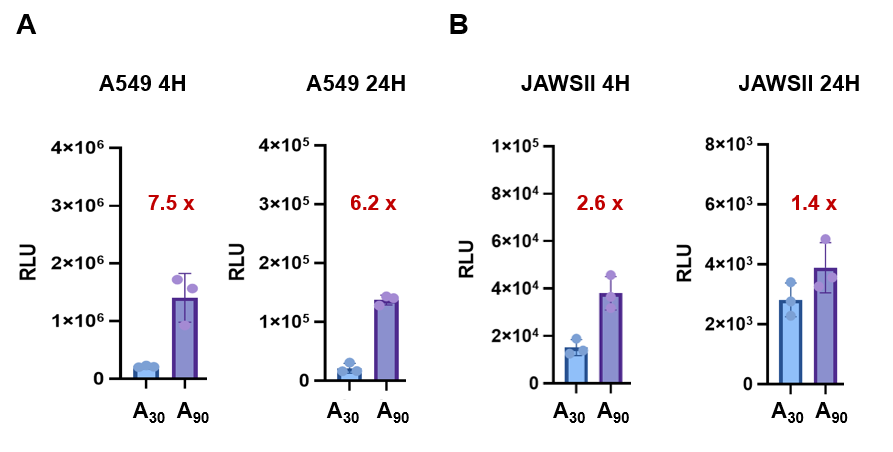

### **Figure S7** **Comparison of FLuc protein expression levels *in vitro* from mRNAs bearing short (A_30_) or long (A_90_) polyA tails.** A) FLuc expression in A549 cells at 4 and 24 hours post-transfection. A substantial increase in expression was observed for the A_90_ variant, as indicated in red. B) FLuc expression in JAWSII cells at 4 and 24 hours post-transfection. The A_90_ variant consistently outperformed A₃₀, although the difference was less pronounced than in A549 cells. Fold changes in red denote the relative increase in expression from A_90_ mRNA compared to A_30_.

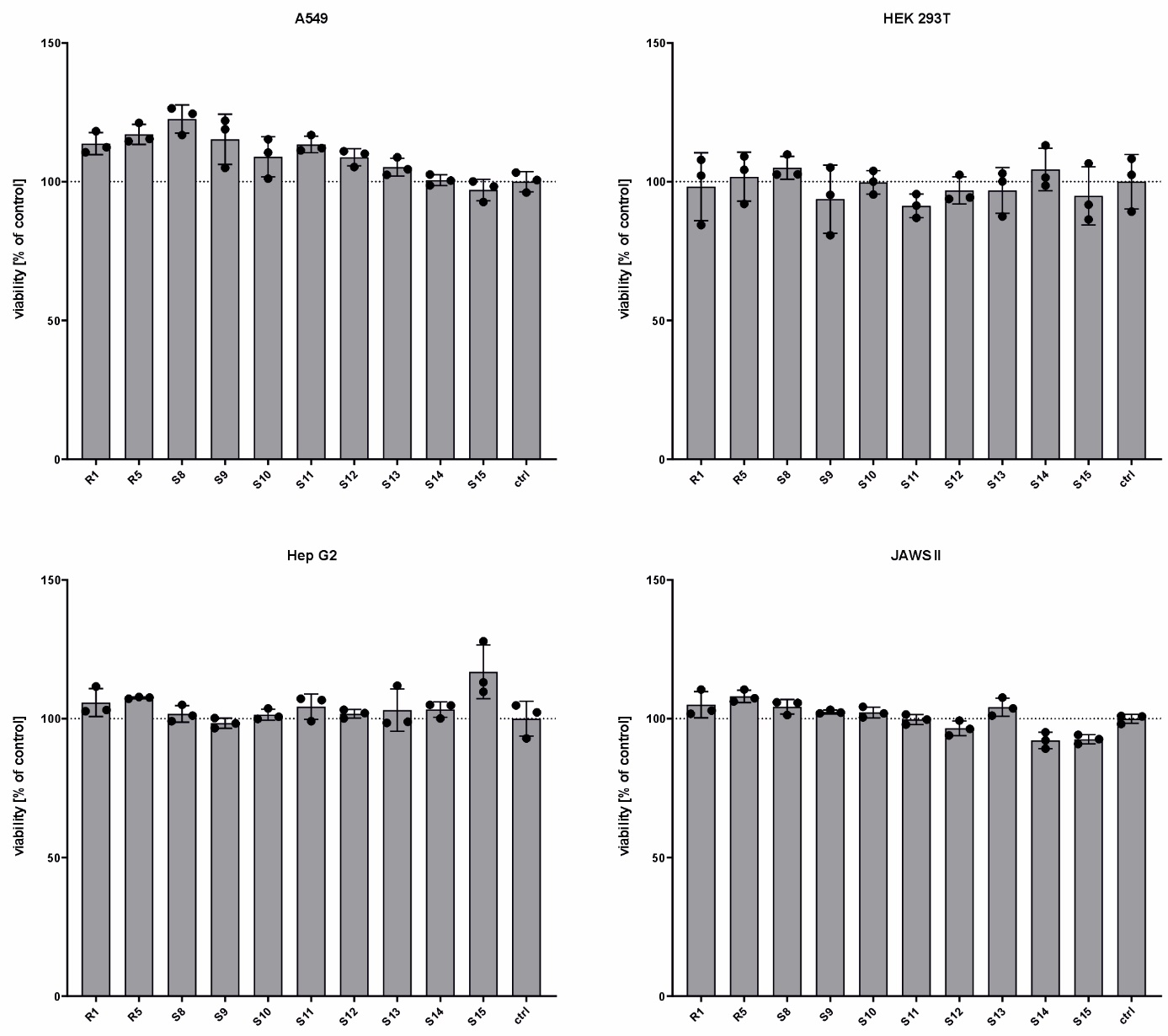

### **Figure S8** **Cell viability following transfection with FLuc mRNAs containing different modified 3’-tail sequences.** Viability of A549, HEK 293T, Hep G2, and JAWS II cells was assessed 24 h post-transfection using Cell Titer Blue assay (Promega). Bars represent viability relative to untreated control cells; means ± SD; n = 3.

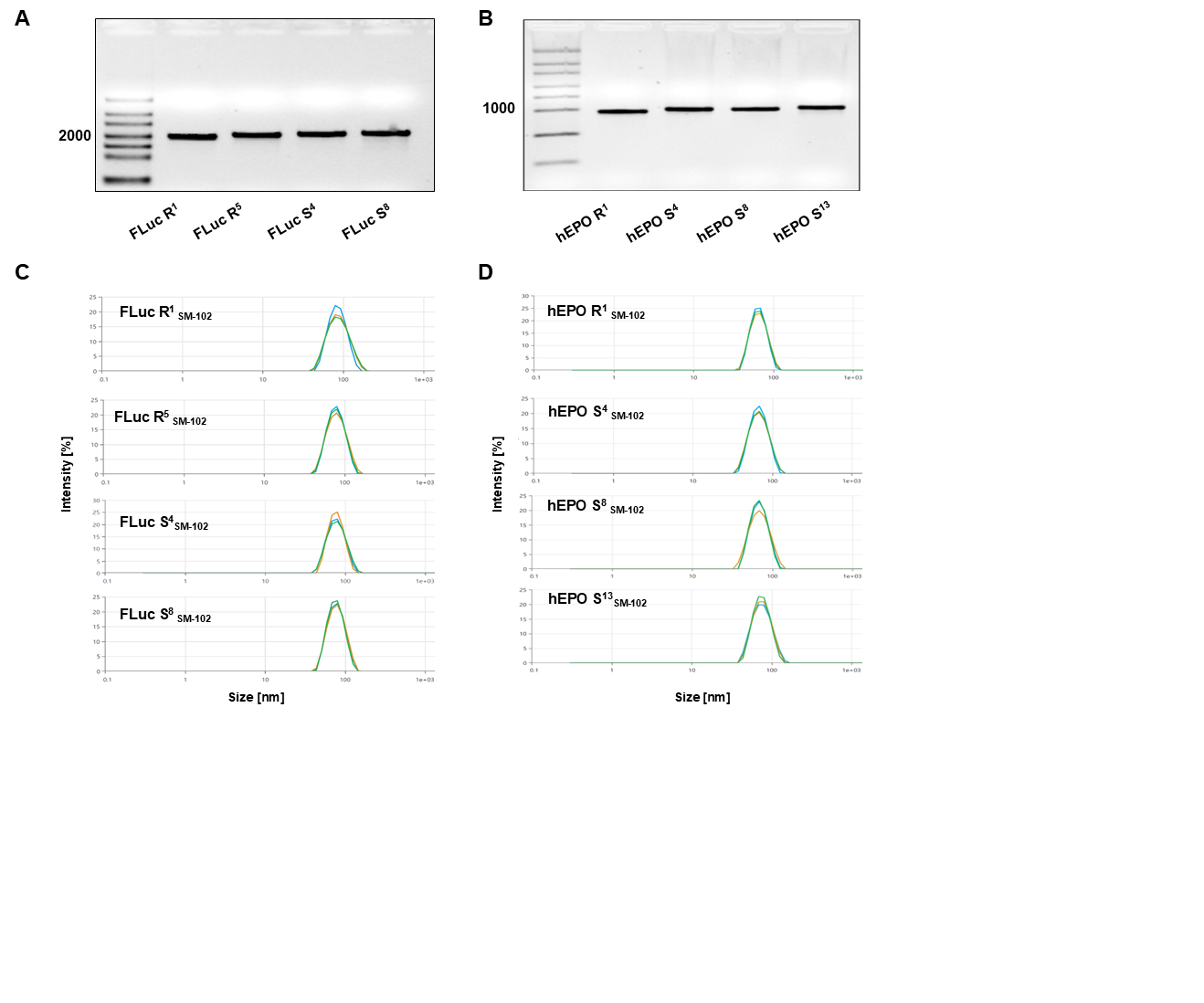

### **Figure S9** **Quality control of lipid nanoparticles preparation for *in vivo* experiments.** A) Electropherograms of all tested mRNA variants, encoding Firefly luciferase (FLuc R^1^ - FLuc S^8^) and B) human erythropoietin (hEPO R^1^ - hEPO S^13^). C) Quality and homogeneity assessment of formulated LNPs (SM-102 lipid mix) for all mRNA variants of FLuc and D) all mRNA variants of hEPO. The graphs illustrate intensity fluctuations over time, determining nanoparticle size and uniformity.

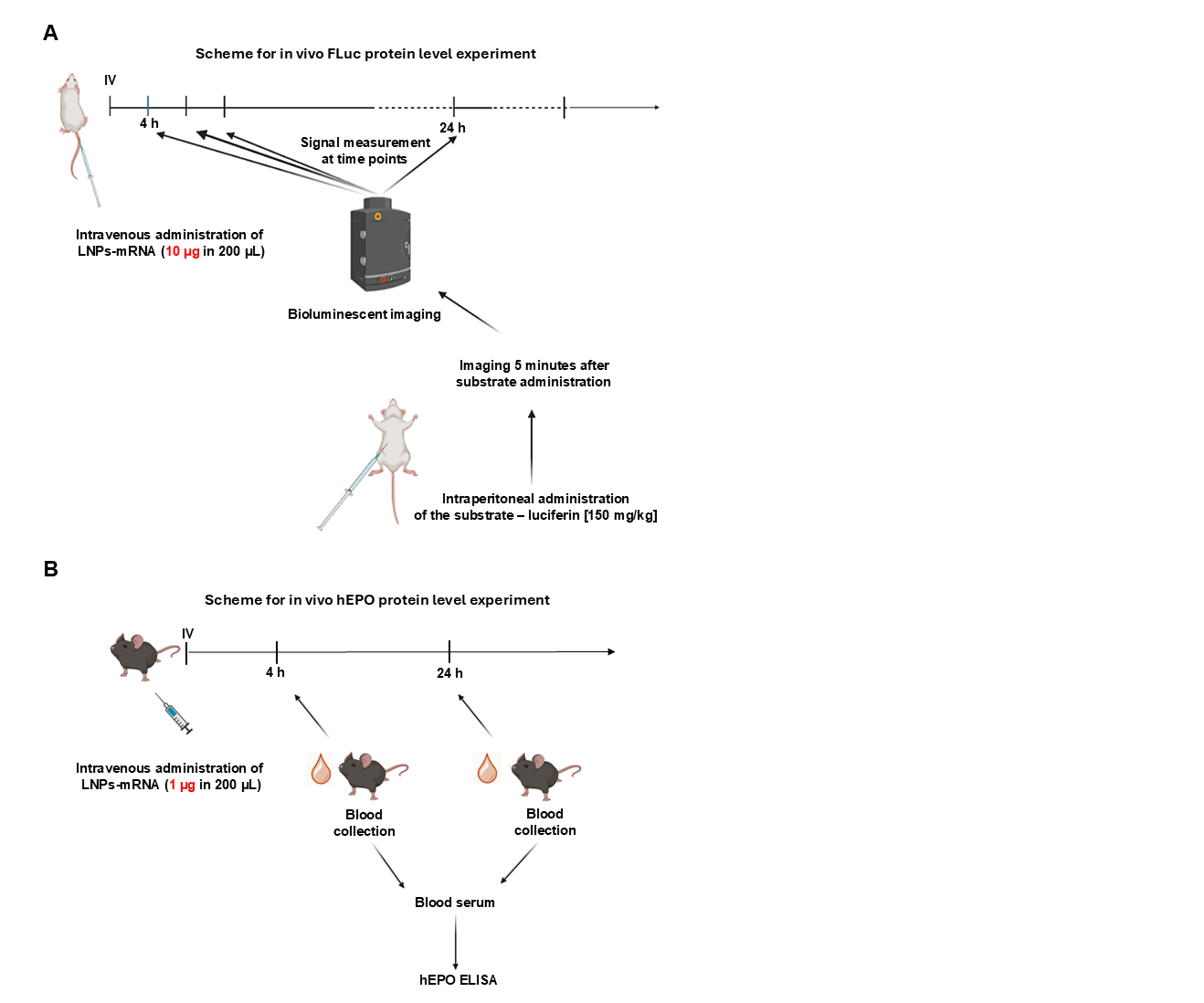

### **Figure S10 Schematic representation of *in vivo* experiments assessing protein expression levels following LNP-mRNA administration.** A) Assessment of *in vivo* FLuc protein expression using bioluminescence imaging. To evaluate the expression of Firefly luciferase (FLuc) *in vivo*, mice received an intravenous (IV) injection of LNP-mRNA (10 µg in 200 µL). Bioluminescence imaging was performed at multiple time points, including 4 h and 24 h post-injection. Five minutes before imaging, luciferin substrate (150 mg/kg) was administered intraperitoneally (IP) to enable bioluminescent signal detection. B) Evaluation of *in vivo* hEPO protein levels through blood serum analysis. To assess the expression of human erythropoietin (hEPO) *in vivo*, mice received an IV injection of LNP-mRNA (1 µg in 200 µL). Blood samples were collected at 4 h and 24 h post-injection to monitor protein levels over time. Serum was then isolated for hEPO quantification using an ELISA assay.

| *Table S1. Summary of complementary DNA oligonucleotide sequences used for the preparation of DNA plasmid templates with modified polyA sequences* | | |
| --- | --- | --- |
| **Mod** | **DNA oligonucleotide sequence** | |
| **R^1^** | Starter F | AAAAAAAAAAAAAAAAAAAAAAAAAAAAAAAAAAAAAAAAAAAAAAAAAAAAAAAAAAAAAA |
|  | Starter R | TTTTTTTTTTTTTTTTTTTTTTTTTTTTTTTTTTTTTTTTTTTTTTTTTTTTTTTTTTTTTT |
| **R^2^** | Starter F | AAAAAAAAAAAAAAAAAAAAAAAAAAAAAAAAGCATATGACTAAAAAAAAAAAAAAAAAAAAAAAAAAAAAAAAAAAAAAAAAAAAAAAAAAAAAAAAAAAAAAAAAAAAAAAA |
|  | Starter R | TTTTTTTTTTTTTTTTTTTTTTTTTTTTTTTTTTTTTTTTTTTTTTTTTTTTTTTTTTTTTTTTTTTTTTTTAGTCATATGCTTTTTTTTTTTTTTTTTTTTTTTTTTTTTTTT |
| **R^3^** | Starter F | AAAAAAAAAAAAAAAAAAAAAAAAAAAAAAAAAAAAAAAAAAAAAAAAAAAAAAAAATCTAG |
|  | Starter R | CTAGATTTTTTTTTTTTTTTTTTTTTTTTTTTTTTTTTTTTTTTTTTTTTTTTTTTTTTTTT |
| **R^4^** | Starter F | AAAAAAAAAAAAAAAAAAAAAAAAAAAAAAAAGAAAAAAAAAAAAAAAAAAAAAAAAAAAAAAAAAAAAAAAAAAAAAAAAAAAAAAAAAAAA |
|  | Starter R | TTTTTTTTTTTTTTTTTTTTTTTTTTTTTTTTTTTTTTTTTTTTTTTTTTTTTTTTTTTTCTTTTTTTTTTTTTTTTTTTTTTTTTTTTTTTT |
| **R^5^** | Starter F | AAAAAAAAAAAAAAAAAAAAAAAAAAAAAAAACAAAAAAAAAAAAAAAAAAAAAAAAAAAAAAAAAAAAAAAAAAAAAAAAAAAAAAAAAAAA |
|  | Starter R | TTTTTTTTTTTTTTTTTTTTTTTTTTTTTTTTTTTTTTTTTTTTTTTTTTTTTTTTTTTTGTTTTTTTTTTTTTTTTTTTTTTTTTTTTTTT |
| **R^6^** | Starter F | AAGAAAAAAAAAAAAAAAAAAAAAAAAAAAAAATGGGGTT TGGGGTTTGGGGTTTGGGGTAAAAAAAAAAAAAAA |
|  | Starter R | TTTTTTTTTTTTTTTACCCCAAACCCCAAACCCCAAACCCCATTTTTTTTTTTTTTTTTTTTTTTTTTTTTTCTT |
| **S^1^** | Starter F | AAGAAAAAAAAAAAAAAAAAAAAAAAAAAAAAAGAAAAAAAAAAAAAAAAAAAAAAAAAAAAAAGAAAAAAAAAAAAAAAAAAAAAAAAAAAAAA |
|  | Starter R | TTTTTTTTTTTTTTTTTTTTTTTTTTTTTTCTTTTTTTTTTTTTTTTTTTTTTTTTTTTTTCTTTTTTTTTTTTTTTTTTTTTTTTTTTTTTCTT |
| **S^2^** | Starter F | AAGCATATAAAAAAAAAAAAAAAAAAAAAAAAAAAAAAGCATATAAAAAAAAAAAAAAAAAAAAAAAAAAAAAAGCATATAAAAAAAAAAAAAAAAAAAAAAAAAAAAAAGCATA TAAAAAAAAAAAAAAAAAAAAAAAAAAAAAA |
|  | Starter R | TTTTTTTTTTTTTTTTTTTTTTTTTTTTTTATATGCTTTTTTTTTTTTTTTTTTTTTTTTTTTTTTATATGCTTTTTTTTTTTTTTTTTTTTTTTTTTTTTTATATGCTTTTTTTTTTTTTTTTTTTTTTTTTTTTTTATATGCTT |
| **S^3^** | Starter F | AAGCATATAAAAAAAAAAAAAAAAAAAAAAAAAAAAAAGCATATAAAAAAAAAAAAAAAAAAAAAAAAAAAAAAGCATATAAAAAAAAAAAAAAAAAAAAAAAAAAAAAA |
|  | Starter R | TTTTTTTTTTTTTTTTTTTTTTTTTTTTTTATATGCTTTTTTTTTTTTTTTTTTTTTTTTTTTTTTATATGCTTTTTTTTTTTTTTTTTTTTTTTTTTTTTTATATGCTT |
| **S^4^** | Starter F | AACAAAAAAAAAAAAAAAAAAAAAAAAAAAAAACAAAAAAAAAAAAAAAAAAAAAAAAAAAAAACAAAAAAAAAAAAAAAAAAAAAAAAAAAAAA |
|  | Starter R | TTTTTTTTTTTTTTTTTTTTTTTTTTTTTTGTTTTTTTTTTTTTTTTTTTTTTTTTTTTTTGTTTTTTTTTTTTTTTTTTTTTTTTTTTTTTGTT |
| **S^5^** | Starter F | AAGAAAAAAAAAAAAAAAGAAAAAAAAAAAAAAAGAAAAAAAAAAAAAAAGAAAAAAAAAAAAAAAGAAAAAAAAAAAAAAAGAAAAAAAAAAAAAAA |
|  | Starter R | TTTTTTTTTTTTTTTCTTTTTTTTTTTTTTTCTTTTTTTTTTTTTTTCTTTTTTTTTTTTTTTCTTTTTTTTTTTTTTTCTTTTTTTTTTTTTTTCTT |
| **S^6^** | Starter F | AAGAAAAAAAAAAAAAAAAAAAAGAAAAAAAAAAAAAAAA AAAAGAAAAAAAAAAAAAAAAAAAAGAAAAAAAAAAAAAA AAAAAAGAAAAAAAAAAAAAAAAAAAA |
|  | Starter R | TTTTTTTTTTTTTTTTTTTTCTTTTTTTTTTTTTTTTTTTTCTTTTTTTTTTTTTTTTTTTTCTTTTTTTTTTTTTTTTTTTTCTTTTTTTTTTTTTTTTTTTTCTT |
| **S^7^** | Starter F | AAGAAAAAAAAAAAAAAAAAAAAAAAAAAAAAAAAAAAAAAAAAAAAAGAAAAAAAAAAAAAAAAAAAAAAAAAAAAAAAAAAAAAAAAAAAAA |
|  | Starter R | TTTTTTTTTTTTTTTTTTTTTTTTTTTTTTTTTTTTTTTTTTTTTCTTTTTTTTTTTTTTTTTTTTTTTTTTTTTTTTTTTTTTTTTTTTTCTT |
| **S^8^** | Starter F | AACAAAAAAAAAAAAAAACAAAAAAAAAAAAAAACAAAAAAAAAAAAAAACAAAAAAAAAAAAAAACAAAAAAAAAAAAAAACAAAAAAAAAAAAAAA |
|  | Starter R | TTTTTTTTTTTTTTTGTTTTTTTTTTTTTTTGTTTTTTTTTTTTTTTGTTTTTTTTTTTTTTTGTTTTTTTTTTTTTTTGTTTTTTTTTTTTTTTGTT |
| **S^9^** | Starter F | TAAAAAAAAAACAAAAAAAAAACAAAAAAAAAACAAAAAAAAAACAAAAAAAAAACAAAAAAAAAACAAAAAAAAAACAAAAAAAAAACAAAAAAAAAACAAAAAAAAAACAAAAAAAAAACAAAAAAAAAA |
|  | Starter R | TTTTTTTTTTGTTTTTTTTTTGTTTTTTTTTTGTTTTTTTTTTGTTTTTTTTTTGTTTTTTTTTTGTTTTTTTTTTGTTTTTTTTTTGTTTTTTTTTTGTTTTTTTTTTGTTTTTTTTTTGTTTTTTTTTTA |
| **S^10^** | Starter F | TAAAAAAAAAAAAACAAAAAAAAAAAAACAAAAAAAAAAAAACAAAAAAAAAAAAACAAAAAAAAAAAAACAAAAAAAAAAAAACAAAAAAAAAAAAACAAAAAAAAAAAAACAAAAAAAAAAAAA |
|  | Starter R | TTTTTTTTTTTTTGTTTTTTTTTTTTTGTTTTTTTTTTTTTGTTTTTTTTTTTTTGTTTTTTTTTTTTTGTTTTTTTTTTTTTGTTTTTTTTTTTTTGTTTTTTTTTTTTTGTTTTTTTTTTTTTA |
| **S^11^** | Starter F | TAAAAAAAAAAAAAAACAAAAAAAAAAAAAAACAAAAAAAAAAAAAAACAAAAAAAAAAAAAAACAAAAAAAAAAAAAAACAAAAAAAAAAAAAAACAAAAAAAAAAAAAAACAAAAAAAAAAAAAAA |
|  | Starter R | TTTTTTTTTTTTTTTGTTTTTTTTTTTTTTTGTTTTTTTTTTTTTTTGTTTTTTTTTTTTTTTGTTTTTTTTTTTTTTTGTTTTTTTTTTTTTTTGTTTTTTTTTTTTTTTGTTTTTTTTTTTTTTTA |
| **S^12^** | Starter F | TAAAAAAAAAAAAAAAAACAAAAAAAAAAAAAAAAACAAAAAAAAAAAAAAAAACAAAAAAAAAAAAAAAAACAAAAAAAAAAAAAAAAACAAAAAAAAAAAAAAAAACAAAAAAAAAAAAAAAAA |
|  | Starter R | TTTTTTTTTTTTTTTTTGTTTTTTTTTTTTTTTTTGTTTTTTTTTTTTTTTTTGTTTTTTTTTTTTTTTTTGTTTTTTTTTTTTTTTTTGTTTTTTTTTTTTTTTTTGTTTTTTTTTTTTTTTTTA |
| **S^13^** | Starter F | TAAAAAAAAAAAAAAAAAAAACAAAAAAAAAAAAAAAAAAAACAAAAAAAAAAAAAAAAAAAACAAAAAAAAAAAAAAAAAAAACAAAAAAAAAAAAAAAAAAAACAAAAAAAAAAAAAAAAAAAA |
|  | Starter R | TTTTTTTTTTTTTTTTTTTTGTTTTTTTTTTTTTTTTTTTTGTTTTTTTTTTTTTTTTTTTTGTTTTTTTTTTTTTTTTTTTTGTTTTTTTTTTTTTTTTTTTTGTTTTTTTTTTTTTTTTTTTTA |
| **S^14^** | Starter F | AACAAAAAAAAAAAAAAACAAAAAAAAAAAAAAACAAAAAAAAAAAAAAACAAAAAAAAAAAAAAACAAAAAAAAAAAAAAACAAAAAAAAAAAAAAACAAAAAAAAAAAAAAACAAAAAAAAAAAAAAACAAAAAAAAAAAAAAACAAAAAAAAAAAAAAA |
|  | Starter R | TTTTTTTTTTTTTTTGTTTTTTTTTTTTTTTGTTTTTTTTTTTTTTTGTTTTTTTTTTTTTTTGTTTTTTTTTTTTTTTGTTTTTTTTTTTTTTTGTTTTTTTTTTTTTTTGTTTTTTTTTTTTTTTGTTTTTTTTTTTTTTTGTTTTTTTTTTTTTTTGTT |
| **S^15^** | Starter F | AACAAAAAAAAAAAAAAAGAAAAAAAAAAAAAAACAAAAAAAAAAAAAAAGAAAAAAAAAAAAAAACAAAAAAAAAAAAAAAGAAAAAAAAAAAAAAACAAAAAAAAAAAAAAAGAAAAAAAAAAAAAAACAAAAAAAAAAAAAAAGAAAAAAAAAAAAAAA |
|  | Starter R | TTTTTTTTTTTTTTTCTTTTTTTTTTTTTTTGTTTTTTTTTTTTTTTCTTTTTTTTTTTTTTTGTTTTTTTTTTTTTTTCTTTTTTTTTTTTTTTGTTTTTTTTTTTTTTTCTTTTTTTTTTTTTTTGTTTTTTTTTTTTTTTCTTTTTTTTTTTTTTTGTT |

| *Table S2. Coding DNA sequences for the Firefly luciferase, mKate2_PEST, human erythropoietin proteins, and* *human alpha-1 antitrypsin used as reporter genes in mRNA studies with modified 3'-tail sequences.* | |
| --- | --- |
| **DNA Sequence Name** | **DNA Sequence** |
| Firefly Lucyferase (FLuc) | ATGGAAGACGCCAAAAACATAAAGAAAGGCCCGGCGCCATTCTATCCTCTAGAGGATGGAACCGCTGGAGAGCAACTGCATAAGGCTATGAAGAGATACGCCCTGGTTCCTGGAACAATTGCTTTTACAGATGCACATATCGAGGTGAACATCACGTACGCGGAATACTTCGAAATGTCCGTTCGGTTGGCAGAAGCTATGAAACGATATGGGCTGAATACAAATCACAGAATCGTCGTATGCAGTGAAAACTCTCTTCAATTCTTTATGCCGGTGTTGGGCGCGTTATTTATCGGAGTTGCAGTTGCGCCCGCGAACGACATTTATAATGAACGTGAATTGCTCAACAGTATGAACATTTCGCAGCCTACCGTAGTGTTTGTTTCCAAAAAGGGGTTGCAAAAAATTTTGAACGTGCAAAAAAAATTACCAATAATCCAGAAAATTATTATCATGGATTCTAAAACGGATTACCAGGGATTTCAGTCGATGTACACGTTCGTCACATCTCATCTACCTCCCGGTTTTAATGAATACGATTTTGTACCAGAGTCCTTTGATCGTGACAAAACAATTGCACTGATAATGAATTCCTCTGGATCTACTGGGTTACCTAAGGGTGTGGCCCTTCCGCATAGAACTGCCTGCGTCAGATTCTCGCATGCCAGAGATCCTATTTTTGGCAATCAAATCATTCCGGATACTGCGATTTTAAGTGTTGTTCCATTCCATCACGGTTTTGGAATGTTTACTACACTCGGATATTTGATATGTGGATTTCGAGTCGTCTTAATGTATAGATTTGAAGAAGAGCTGTTTTTACGATCCCTTCAGGATTACAAAATTCAAAGTGCGTTGCTAGTACCAACCCTATTTTCATTCTTCGCCAAAAGCACTCTGATTGACAAATACGATTTATCTAATTTACACGAAATTGCTTCTGGGGGCGCACCTCTTTCGAAAGAAGTCGGGGAAGCGGTTGCAAAACGCTTCCATCTTCCAGGGATACGACAAGGATATGGGCTCACTGAGACTACATCAGCTATTCTGATTACACCCGAGGGGGATGATAAACCGGGCGCGGTCGGTAAAGTTGTTCCATTTTTTGAAGCGAAGGTTGTGGATCTGGATACCGGGAAAACGCTGGGCGTTAATCAGAGAGGCGAATTATGTGTCAGAGGACCTATGATTATGTCCGGTTATGTAAACAATCCGGAAGCGACCAACGCCTTGATTGACAAGGATGGATGGCTACATTCTGGAGACATAGCTTACTGGGACGAAGACGAACACTTCTTCATAGTTGACCGCTTGAAGTCTTTAATTAAATACAAAGGATATCAGGTGGCCCCCGCTGAATTGGAATCGATATTGTTACAACACCCCAACATCTTCGACGCGGGCGTGGCAGGTCTTCCCGACGATGACGCCGGTGAACTTCCCGCCGCCGTTGTTGTTTTGGAGCACGGAAAGACGATGACGGAAAAAGAGATCGTGGATTACGTCGCCAGTCAAGTAACAACCGCGAAAAAGTTGCGCGGAGGAGTTGTGTTTGTGGACGAAGTACCGAAAGGTCTTACCGGAAAACTCGACGCAAGAAAAATCAGAGAGATCCTCATAAAGGCCAAGAAGGGCGGAAAGTCCAAATTGTAA |
| mKate2-PEST | ATGGTCAGCGAGTTGATCAAGGAGAACATGCACATGAAGCTGTATATGGAGGGGACCGTCAACAACCACCACTTCAAATGCACGTCCGAGGGGGAGGGCAAACCGTACGAGGGCACCCAGACGATGCGGATCAAGGCCGTCGAGGGGGGCCCtCTCCCCTTCGCCTTCGATATCCTCGCGACCAGCTTCATGTACGGCTCCAAGACGTTCATCAACCACACGCAGGGCATCCCAGACTTCTTCAAGCAGTCGTTCCCGGAAGGGTTCACGTGGGAGAGGGTGACCACGTACGAAGACGGTGGGGTGTTGACCGCTACGCAAGACACGTCCCTCCAGGACGGCTGCCTGATTTACAACGTCAAGATCCGGGGCGTCAACTTCCCGAGCAACGGGCCCGTAATGCAGAAGAAGACTTTGGGGTGGGAGGCCTCGACCGAGACCTTGTACCCCGCCGACGGCGGCCTTGAGGGGCGAGCTGACATGGCTCTCAAGCTCGTCGGCGGGGGACACTTGATCTGCAACCTAAAAACGACGTACAGGTCCAAGAAGCCGGCGAAGAACCTAAAGATGCCTGGCGTCTACTACGTGGACCGGAGGCTCGAGAGGATCAAGGAGGCGGACAAAGAGACCTACGTGGAGCAGCACGAGGTGGCAGTCGCCCGCTACTGCGATCTCCCCAGTAAGCTCGGCCACCGGAGCCACGGATTCCCTCCAGCAGTTGCTGCTCAAGACGACGGGACTCTGCCCATGTCCTGTGCGCAAGAATCTGGAATGGATCGACATCCTGCAGCTTGCGCCAGCGCTAGAATCAACGTGTAA |
| Human erythropoietin (hEPO) | ATGGGCGTGCACGAGTGCCCCGCCTGGCTGTGGCTGCTGCTGAGCCTGCTGAGCCTGCCCCTGGGCCTGCCCGTGCTGGGCGCCCCCCCCCGGCTGATCTGCGACAGCCGGGTGCTGGAGCGGTACCTGCTGGAGGCCAAGGAGGCCGAGAACATCACCACCGGCTGCGCCGAGCACTGCAGCCTGAACGAGAACATCACCGTGCCCGACACCAAGGTGAACTTCTACGCCTGGAAGCGGATGGAGGTGGGCCAGCAGGCCGTGGAGGTGTGGCAGGGCCTGGCCCTGCTGAGCGAGGCCGTGCTGCGGGGCCAGGCCCTGCTGGTGAACAGCAGCCAGCCCTGGGAGCCCCTGCAGCTGCACGTGGACAAGGCCGTGAGCGGCCTGCGGAGCCTGACCACCCTGCTGCGGGCCCTGGGCGCCCAGAAGGAGGCCATCAGCCCCCCCGACGCCGCCAGCGCCGCCCCCCTGCGGACCATCACCGCCGACACCTTCCGGAAGCTGTTCCGGGTGTACAGCAACTTCCTGCGGGGCAAGCTGAAGCTGTACACCGGCGAGGCCTGCCGGACCGGCGACCGGTGA |
| Human alpha-1 antitrypsin  (hA1AT) | ATGCCTAGCAGCGTGTCATGGGGTATTCTGCTGCTGGCCGGACTGTGTTGTCTGGTGCCTGTGTCTCTGGCTGAAGATCCTCAAGGCGACGCCGCTCAGAAAACCGATACAAGCCACCACGACCAGGATCACCCCACCTTCAACAAGATCACCCCTAACCTGGCCGAGTTCGCCTTCAGCCTGTATAGACAGCTGGCCCACCAGAGCAACAGCACCAACATCTTTTTCAGCCCCGTGTCTATCGCCACCGCCTTTGCTATGCTGAGCCTGGGCACAAAGGCCGACACACACGATGAGATCCTGGAAGGCCTGAACTTCAACCTGACAGAGATCCCCGAGGCTCAGATCCACGAGGGCTTTCAAGAGCTGCTGAGAACCCTGAACCAGCCTGACTCTCAGCTCCAGCTGACAACCGGCAATGGCCTGTTTCTGTCTGAGGGCCTGAAGCTGGTGGACAAGTTCCTGGAAGATGTGAAGAAGCTGTACCACAGCGAGGCCTTCACCGTGAACTTCGGCGATACCGAGGAAGCCAAGAAGCAGATCAACGACTACGTGGAAAAGGGCACCCAGGGCAAGATCGTGGACCTGGTCAAAGAGCTGGACAGAGACACCGTGTTCGCCCTGGTCAACTACATCTTCTTCAAAGGCAAGTGGGAACGCCCCTTCGAAGTGAAGGACACAGAGGAAGAGGACTTCCACGTCGACCAAGTGACCACCGTGAAGGTGCCCATGATGAAGCGGCTGGGCATGTTCAACATCCAGCACTGCAAGAAACTGAGCAGCTGGGTGCTGCTGATGAAGTACCTGGGCAACGCCACAGCCATATTCTTTCTGCCCGATGAGGGCAAGCTGCAGCACCTGGAAAATGAGCTGACCCACGACATCATCACCAAGTTCCTAGAGAACGAGGACAGAAGAAGCGCCAGCCTGCATCTGCCTAAGCTGAGCATCACCGGCACCTACGATCTGAAGTCTGTGCTGGGACAGCTGGGCATCACAAAGGTGTTCAGCAATGGCGCCGATCTGTCCGGCGTTACAGAAGAGGCTCCTCTGAAGCTGTCCAAGGCCGTGCACAAAGCCGTGCTGACAATCGATGAGAAGGGAACAGAGGCCGCTGGCGCCATGTTTCTGGAAGCTATCCCTATGAGCATCCCGCCTGAAGTGAAGTTCAACAAGCCCTTCGTGTTCCTGATGATCGAACAGAATACCAAGTCTCCCCTGTTCATGGGCAAAGTGGTCAACCCCACACAGAAATGA |

| *Table S3. mRNA half-lives and translational efficiencies for mKate2_PEST mRNA with modified 3’-tail sequences for A549 and HEK293T cell lines.* | | | | | |
| --- | --- | --- | --- | --- | --- |
| **Cell line** | **mRNA variant** | **mRNA half-life [min]** | **mRNA half-life SD [min]** | **mRNA translation rate** | **mRNA translation rate SD** |
| A549 | mKate2_R^1^ | 348,93 | 15,50 | 0,39 | 0,04 |
|  | mKate2_R^2^ | 321,49 | 7,02 | 0,47 | 0,08 |
|  | mKate2_R^3^ | 406,74 | 32,07 | 0,55 | 0,06 |
|  | mKate2_R^4^ | 230,43 | 17,21 | 0,34 | 0,08 |
|  | mKate2_R^5^ | 349,32 | 66,68 | 0,54 | 0,10 |
|  | mKate2_R^6^ | 41,59 | 0,00 | 0,01 | 0,00 |
|  | mKate2_S^1^ | 415,08 | 35,41 | 0,59 | 0,07 |
|  | mKate2_S^2^ | 416,83 | 40,19 | 0,41 | 0,06 |
|  | mKate2_S^3^ | 347,74 | 14,74 | 0,36 | 0,06 |
|  | mKate2_S^4^ | 417,24 | 13,09 | 0,65 | 0,03 |
|  | mKate2_S^5^ | 426,73 | 8,97 | 0,71 | 0,09 |
|  | mKate2_S^6^ | 455,27 | 28,46 | 0,88 | 0,11 |
|  | mKate2_S^7^ | 380,67 | 33,04 | 0,65 | 0,06 |
|  | mKate2_S^8^ | 413,14 | 28,47 | 0,77 | 0,10 |
| 293T | mKate2_R^1^ | 342,21 | 24,99 | 0,97 | 0,08 |
|  | mKate2_R^2^ | 345,30 | 55,08 | 1,09 | 0,04 |
|  | mKate2_R^3^ | 441,80 | 64,57 | 1,19 | 0,13 |
|  | mKate2_R^4^ | 248,10 | 20,51 | 0,68 | 0,10 |
|  | mKate2_R^5^ | 399,63 | 15,29 | 1,05 | 0,32 |
|  | mKate2_R^6^ | 41,59 | 0,00 | 0,01 | 0,01 |
|  | mKate2_S^1^ | 363,04 | 13,93 | 1,22 | 0,15 |
|  | mKate2_S^2^ | 339,49 | 8,07 | 0,88 | 0,13 |
|  | mKate2_S^3^ | 254,31 | 21,35 | 0,85 | 0,12 |
|  | mKate2_S^4^ | 298,53 | 14,23 | 1,15 | 0,10 |
|  | mKate2_S^5^ | 296,63 | 17,42 | 1,24 | 0,01 |
|  | mKate2_S^6^ | 343,43 | 33,19 | 1,39 | 0,17 |
|  | mKate2_S^7^ | 316,60 | 17,97 | 1,19 | 0,15 |
|  | mKate2_S^8^ | 303,56 | 22,67 | 1,15 | 0,12 |
